# Virus-encoded glycosyltransferases hypermodify DNA with diverse glycans

**DOI:** 10.1101/2023.12.21.572611

**Authors:** Jesse D. Pyle, Sean R. Lund, Katherine H. O’Toole, Lana Saleh

**Affiliations:** Research Department, New England Biolabs, 240 County Road, Ipswich, MA 01938, USA

**Keywords:** Bacteriophage, DNA hypermodification, DNA glycans, TET, 5-methylpyrimidine dioxygenase, 5-hydroxymethylcytosine, glycosyltransferase, biosynthetic gene cluster

## Abstract

Enzymatic modification of DNA nucleobases can coordinate gene expression, protection from nucleases, or mutagenesis. We recently discovered a new clade of phage-specific cytosine methyltransferase (MT) and 5-methylpyrimidine dioxygenase (5mYOX, *e.g.,* TET) enzymes that produce 5-hydroxymethylcytosine (5hmC) as a precursor for additional post-replicative enzymatic hypermodifications on viral genomes. Here, we identify phage MT- and 5mYOX-dependent glycosyltransferase (GT) enzymes that catalyze linkage of diverse glycans directly onto 5hmC reactive nucleobase substrates. Using targeted bioinformatic mining of the phage metavirome databases, we discovered thousands of new biosynthetic gene clusters (BGCs) containing enzymes with predicted roles in cytosine sugar hypermodification. We developed a pathway reassembly platform for high-throughput functional screening of GT-containing BGCs, relying on the endogenous *E. coli* metabolome as a substrate pool. We successfully reconstituted a subset of phage BGCs and isolated novel and highly diverse sugar modifications appended to 5hmC, including mono-, di-, or tri-saccharide moieties comprised of hexose, N-acetylhexosamine or heptose sugars. Structural predictions and sugar product analyses suggest that phage GTs are related to host lipopolysaccharide, teichoic acid, and other small molecule biosynthesis enzymes and have been repurposed for DNA substrates. An expanded metagenomic search revealed hypermodification BGCs within gene neighborhoods containing phage structural proteins and putative genome defense systems. These findings enrich our knowledge of secondary modifications on DNA and the origins of corresponding sugar writer enzymes. Post-replicative cytosine hypermodification by virus-encoded GTs is discussed in the context of genome defense, DNA partitioning and virion assembly, and host-pathogen co-evolution.

## INTRODUCTION

Mature biomolecules are subject to a broad array of enzymatic modifications that regulate structure and function. The physicochemical plasticity afforded by these modifications expands the operational landscape of the central dogma and the responses of biological systems to exogenous and endogenous stimuli (1–3). Epigenetic DNA signatures, post-transcriptional modifications on all subtypes of cellular RNA, and diverse post-translational protein modifications are examples of such variation. Once viewed as a static carrier of genetic information, genomic DNA can serve as a substrate for targeted and dynamic enzymatic chemical modifications. DNA modification by base methylation, for example, occurs across all domains of life and functions in epigenetic control of gene expression, chromatin reorganization, DNA protection and repair (4). Additional DNA modification systems have evolved out of immune defense responses during host-pathogen co-evolution, including restriction-modification systems in prokaryotes (5), antiviral mutagenesis (*e.g.,* APOBEC3 enzymes) and antibody diversification (*e.g.,* AID enzymes) mechanisms in higher eukaryotes (6, 7).

Viruses are the most abundant (8, 9) and genetically diverse (10) agents in the biosphere. Viruses of bacteria (phage) vastly outnumber their ubiquitous host cells and are unappreciated regulators of global nutrient cycling via modulation of host populations (11, 12). The ancient evolutionary arms race between phage and bacteria has yielded sophisticated genome defense strategies that have been adopted and adapted by both host and pathogen. Invading phage genomes are targets for bacterial restriction-modification and CRISPR-Cas systems, which bind and cleave DNA at sequence-specific regions of the viral genome (5, 13, 14). As a result, some phage have evolved antagonistic systems to modify their DNA bases, in-part for steric hinderance of bacterial nucleases (15–17).

In phage DNA, modifications to pyrimidine bases can be installed pre- or post-replication through the enzymatic actions of phage-encoded thymidylate synthase or 5-methylpyrimidine dioxygenase (5mYOX) family enzymes, respectively (16, 18, 19). A well-studied subset of the 5mYOX superfamily involved in DNA base modification is the ten-eleven translocase (TET) family of enzymes. In higher eukaryotes, TET coordinates the recycling of epigenetic markers on cytosine via successive 2-oxoglutarate and iron(II)-dependent oxidation of 5-methylcytosine (5mC) to 5-hydroxymethylcytosine (5hmC), 5-formylcytosine (5fC), and 5-carboxycytosine (5caC) (**Fig. 1A**), the latter two products being subjected to base excision repair and reversion of the intrapolymer base to cytosine (20–23). Phage 5mYOXs, by contrast, have been observed to stall oxidation at 5hmC via an unknown mechanism, allowing for accumulation of a nucleophilic 5hmC hydroxyl handle (**Fig. 1B**) for subsequent chemistries on the DNA base (18). This is thought to be the primary post-replicative mechanism by which phage enzymes install complex chemical modification onto cytosine bases in their genomic DNA.

**Figure 1.**
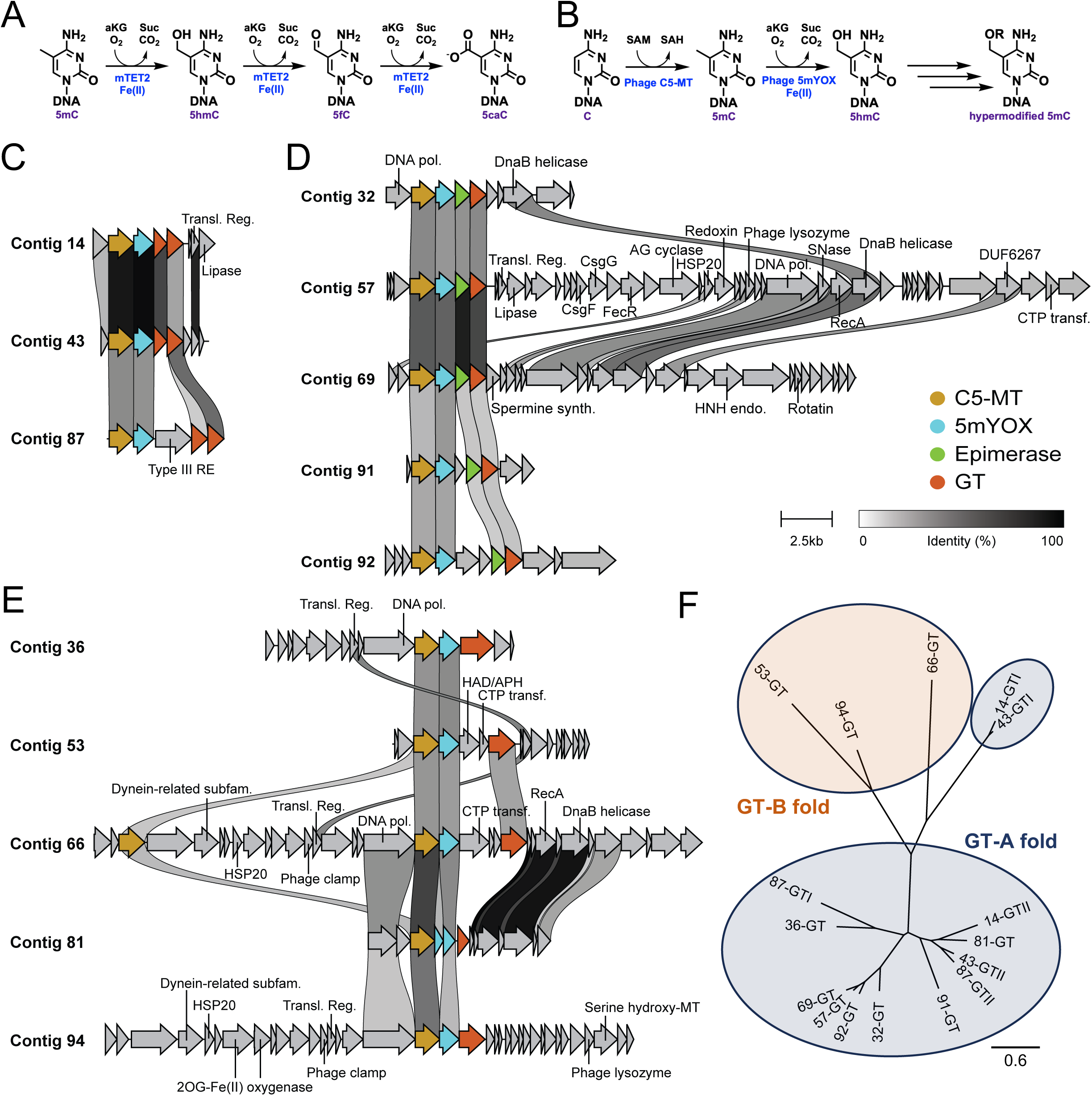
5mYOX-associated GTs with predicted functions in DNA hypermodification encoded by phage genomes. (A) Murine TET2 (mTET2) performs sequential oxidation of 5mC using an Fe(II) and alpha-ketoglutarate (aKG)-dependent reaction mechanism. (B) General pathway of cytosine modification by phage-encoded C5-MT, 5mYOX, and neighboring hypermodification enzymes. (C-E) Synteny visualization for subclass I (C), subclass II (D), and subclass III (E) phage pathways. Subclasses are defined by the presence of sequential GTs (I), a paired sugar epimerase with a neighboring GT (II), or a GT paired with another potential hypermodification-associated enzyme cluster (III). Pathway synteny visualized using clinker software. Scale bars for pathway length and percent identity between coding regions is shown to the right. (F) Phylogenetic analysis of phage-encoded GTs. Ovals represent enzymes with GT-A or GT-B folds as predicted using AlphaFold. Scale bar represents amino acid substitutions per site.

Phage 5mYOXs are encoded in the viral genome within biosynthetic gene cluster (BGC) neighborhoods adjacent to C5-cytosine methyltransferase (C5-MT) enzymes (18), reflecting the stepwise chemistry of the DNA modification pathway (**Fig. 1B**). Unlike eukaryotic DNA methylation patterns, most phage C5-MTs have a GpC sequence context preference (18), providing a level of specificity for subsequent 5mYOX oxidation and 5hmC installation. Another gene family commonly harbored in the same operon as C5-MT and 5mYOX are putative DNA glycosyltransferases (GTs) (18, 24). We have shown that C5-MT/5mYOX-adjacent GTs can catalyze transfer of sugar residues to 5hmC sites in the DNA polymer (18). Some phage have highly glycosylated genomes (25, 26) resulting from pre-replicative manipulation of cellular NTP pools for incorporation of 5hmC at most or all sites in newly synthesized viral genomes (25–28) followed by sugar addition by phage-encoded GTs (29). This widespread incorporation of modified bases can function to protect phage genomes from host nuclease defenses (30–34). The GpC-specific nature of post-replicatively modified DNA (18) suggests alternative modification-dependent functions beyond sugar-coating of the viral genome strictly for defense against restriction enzymes, as R-M systems have broader specificity that could circumvent the protection afforded by these modifications.

Our knowledge of chemical modifications to canonical nucleic acid structures continues to expand, prompting new questions about the nature and plasticity of the genetic “language”. Here, we report the discovery of ten new sugar modifications to cytosine resulting from a limited sample size of recently discovered 5mYOX-associated metavirome DNA GTs. We developed an *E. coli*-based discovery platform for phage BGC reassembly and used this system to screen metavirome pathways, validate enzyme activities, and identify novel DNA hypermodifications. Enzymatic and bioinformatic analyses of phage hypermodification GTs reveal functional overlaps with bacterial enzymes, highlighting a previously underappreciated connection between host-pathogen adaptation and enzyme evolution. Our findings reflect what we believe to be a narrow glimpse of the unexplored chemical possibilities tolerated by DNA in biological systems and suggest that abundant new hypermodifications, and their biological roles, remain to be discovered in the natural world.

## RESULTS

### Discovery of GT-containing gene clusters from phage metaviromes

We previously identified diverse metavirome pathways containing C5-MT and 5mYOX-associated enzymes with predicted sugar binding and modification functions (18). This initial search was limited to contig assemblies contained in early iterations of the GOV2.0 and IMG/VR metagenomic databases (35, 36). Within this dataset of C5-MT and 5mYOX-containing pathways, we identified 21 predicted GTs adjacent to phage 5hmC-synthesizing enzymes (**Fig. 1C-E**). Phylogenetic (**Fig. 1F**) and multiple sequence alignment (MSA, **Fig. S1**) analyses revealed limited sequence similarity among the GTs, suggesting diverse origins or evolutionary pressures for these enzymes. Enzymatic diversity is further supported by differences in computational predictions of structure (*e.g.,* A vs. B form GT) and function (*e.g.,* retaining vs. inverting mechanisms) for the newly identified phage GTs (**Fig. S1, Table 1**).

**Table 1.**
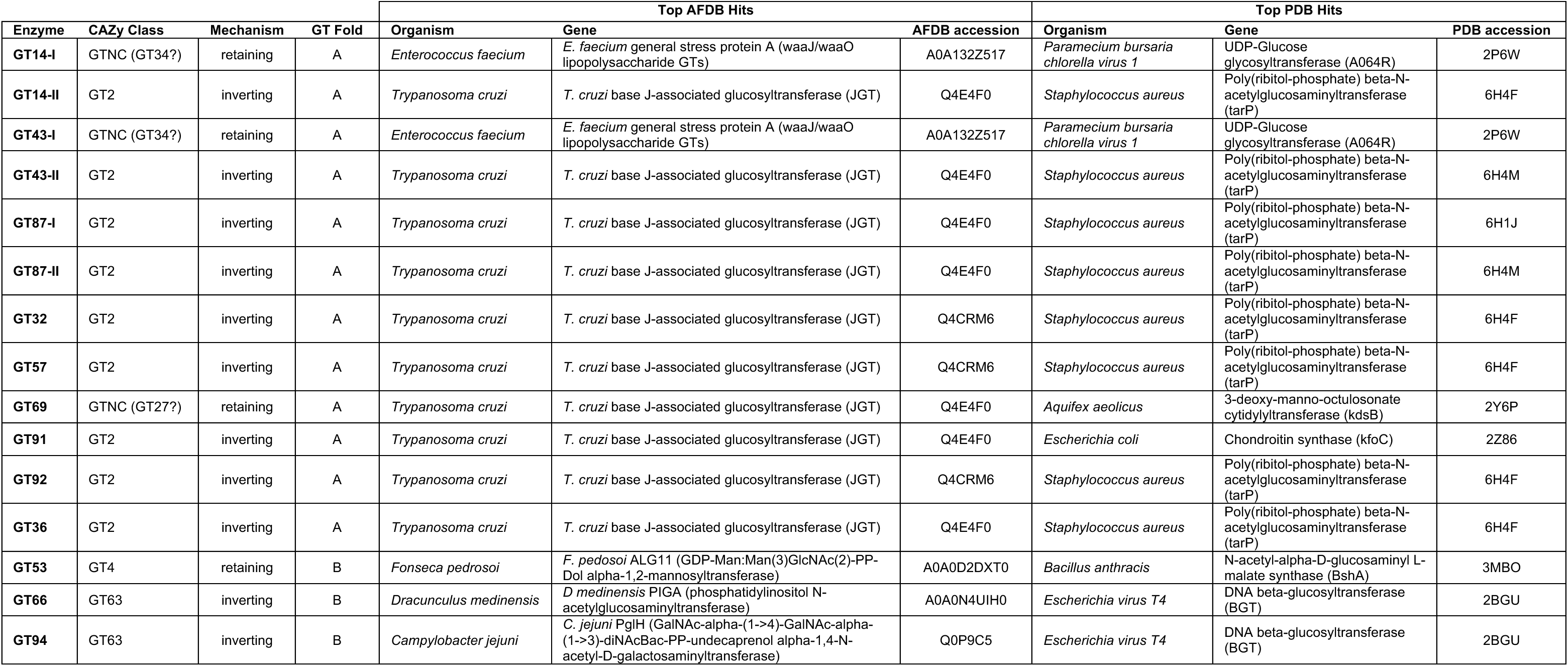
Properties and structural homologs of phage-encoded DNA glycosyltransferases. Structural models for each phage GT were predicted using AlphaFold and used for structure-guided searches of the AlphaFold Proteome Database (AFDB) and Protein Data Bank (PDB) of experimentally determined macromolecular structures. Phage GT sequences were assigned to a CAZy class based on the top structural homologs from the PDB.

Contig synteny revealed three subclasses of GT-containing sequences defined by general shared pathway architecture within each subclass. Subclass I pathways encode two successive GT enzymes neighboring the C5-MT and 5mYOX (**Fig. 1C**). Subclass II pathways share a GT and a sugar epimerase adjacent pairing (**Fig. 1D**). Subclass III includes pathways containing a GT and at least one other predicted hypermodification enzyme, although the most organizational variation is found among contigs within this subclass (**Fig. 1E**). The variation in GT primary sequence does not correlate with contig subclass designation, except for a cluster of epimerase-dependent GT sequences with ∼55-78% identity (**Figs. 1F** **and S1**). The considerable variation in GT sequence and pathway context prompted us to develop a scalable high-throughput (HT) system to probe the biosynthetic capacity of these novel phage pathways and assess their potential functions in cytosine hypermodification.

### BGC reassembly using a HT *in vivo* discovery platform

To address the enzymology and products of novel phage pathways, we developed a BGC reconstitution system functioning with endogenous metabolites of *E. coli*. The assembly design is described in detail within Materials and Methods. We focused on developing a co-expression platform for all enzymes in a novel pathway of interest, and with each enzyme under separate transcription control by T7 RNA polymerase. Briefly, genes of interest with T7 promoter and terminator regions are amplified using position-specific primers, allowing for one-step assembly into a single plasmid expression vector using Golden Gate Assembly (37–40) (**Fig. 2A**). Position of the enzymes and correct construct assembly is verified by colony PCR or whole-plasmid sequencing. Enzyme co-expression has been tested with up to five inserted genes of interest (**Fig. S2**) although this assembly approach allows for dozens of correctly assembled fragments with only moderate losses in assembly efficiency (41). Using liquid handling robotics, this system is used to design and assemble dozens to hundreds of different pathways and enzyme combinations in parallel, including combinatorial and leave-one-out permutations for each individual BGC (**Fig. S2**). Assembled constructs are transformed into *E. coli* with *lac* inducible expression of T7 RNA polymerase to coordinate transcription for the assembled genes of interest.

**Figure 2.**
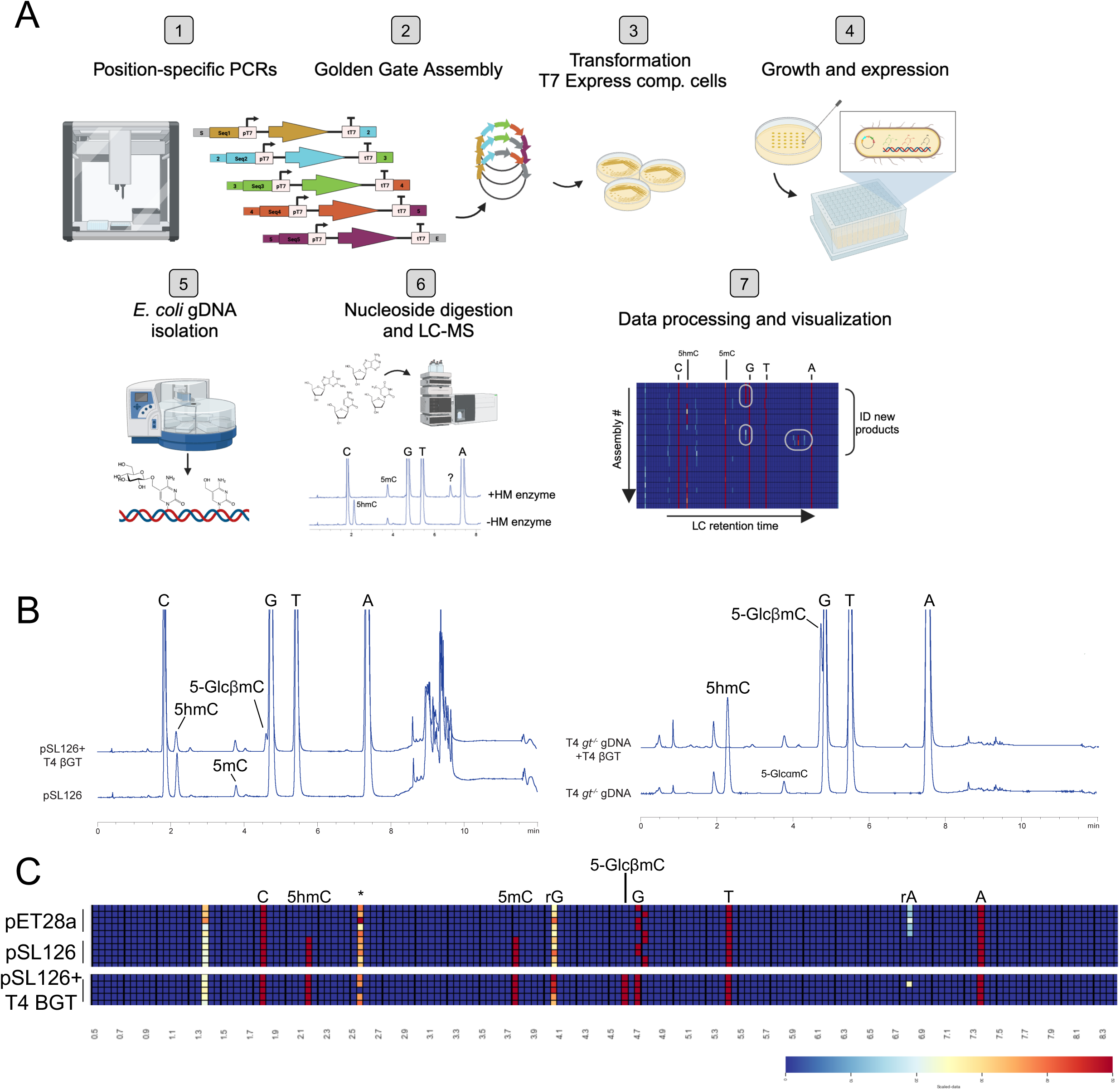
A high-throughput platform for parallel analysis of novel biosynthetic gene clusters. (A) Steps describing the workflow for reconstituting BGCs in *E. coli* and downstream analysis of modified DNA products. (1) Assemblies are designed from a database of metavirome pathway coding regions and genes of interest are amplified by PCR using position-specific primers with the assistance of the Opentrons liquid handling system. (2) Amplicons are assembled into final expression vectors by Golden Gate Assembly. (3) Assembled plasmids are transformed into NEB T7 Express E. coli and individual colonies are picked and grown in 96-well plates for (4) pathway expression by IPTG induction and parallel replica plates for cPCR verification. (5) Modified E. coli gDNA is isolated using a KingFisher automation instrument and subjected to (6) nucleoside digestion and LC-MS analysis. (7) Data from individual assemblies are processed and masses for new products are determined. (B) BGC expression system validation using co-expression of C5-MT and 5mYOX. Left: Plasmid pSL126 carrying MT14 and TET43 was used as the destination GGA vector for T4 βGT, which glucosylates 5hmC precursor bases. Right: Comparison against UHPLC traces from *in vitro* glucosylated DNA. Nucleoside compounds eluting within each UHPLC peak were verified by MS and are labeled. (C) Heat map visualization of peak areas for parallel gDNA samples from *E. coli* expressing empty vector (pET28a), MT14+TET43 alone (pSL126), and MT14+TET43+T4 βGT (pSL126+T4 βGT). Five replicates are shown for each condition. Heat map scale by color intensity is shown below and corresponds to total normalized area for peaks across retention time intervals. The elution of a non-specific product peak of unknown mass is indicated with an asterisk.

This is an expansion of our previously established cell-based pathway analysis system (18), where multiple plasmids were used to co-express phage C5-MT and 5mYOX enzymes in *E. coli*. These intracellular enzymes act on endogenous sources of DNA – the *E. coli* genomic DNA (gDNA) and transformed plasmid DNA – which provides unbiased sequence contexts for C5-MT base methylation, 5mYOX oxidation, and subsequent chemical installations (**Figs. 1B**) (18). The same activity occurs in our BGC assembly platform, where expressed enzymes have access to the endogenous *E. coli* metabolome for suitable substrates and act on the available DNA within the cell (**Fig. 2A**). To detect novel modifications directly on the DNA bases, we isolated the *E. coli* gDNA and digested the polymer to individual nucleosides. The nucleoside pool is analyzed using reverse-phase liquid chromatography mass spectrometry (LC-MS), as described previously (18). Novel products from total digested *E. coli* gDNA typically elute within a 1.5-to-7-minute LC retention time window, among peaks for canonical dC, dG, dT, and dA nucleosides (**Fig. 2B**). Analytes were continuously monitored by mass spectrometry in positive and negative mode during LC separation and adduct ion masses were assigned to novel peaks. For most sugar-modified nucleosides, we were able to detect [M+H]^+^, [M+Na]^+^, and [M+K]^+^ adduct ions in positive mode and [M-H]^-^ and [M+CH_3_CO_2_-H]^-^ adduct ions in negative mode (**Fig. S3**).

We previously identified highly active C5-MT (MT14) and 5mYOX (TET43) that produced the most abundant levels of 5mC and 5hmC in our *in vivo* expression system relative to other metavirome pathways tested (18). As a starting point for screening the activities of novel hypermodification enzymes, we co-expressed MT14 and TET43 in each of our construct designs to provide as much 5hmC substrate as possible and increase the likelihood of modification detection. Single-plasmid expression of MT14 and TET43 produced detectable levels of 5mC and 5hmC on bacterial gDNA (**Fig. 2B**) similar to previous co-expression of these enzymes on separate plasmids (18). Co-expression of β-glucosyltransferase from T4 phage (T4-βGT) in the same plasmid resulted in an additional peak for β-glucosyl-5-methylcytosine (5-GlcβmC) representing conversion of TET43-synthesized 5hmC to hypermodified glucosylated 5mC (**Fig. 2B**). Samples are processed in 96-well plate format and the data visualized as heat maps displaying relative abundance of canonical and modified nucleosides (**Figs. 2C** **and S2-3**). These findings establish and validate our HT BGC platform as a robust system for parallel assessment of novel hypermodification pathway enzyme functions.

### Phage BGC enzymes install novel sugars on cytosine

Systematic *in vivo* reconstitution of GT-containing phage pathways resulted in the production of original and highly diverse sugar-modified cytosine bases (**Fig. 3A-B**). The sugars appended to cytosine appeared to include six-, seven-, or eight-carbon sugars, mono-, di-, or trisaccharide glycans, and ulosonic (*e.g.,* Kdo) and uronic (HexA) sugar acids (**Fig. 3B**). Sugar type generally did not correlate with BGC subclass (**Fig. 3B**). Some sugars with identical masses (*e.g.,* 419 amu products from contigs 81 and 87) had different LC retention times suggesting isomeric or anomeric differences.

**Figure 3.**
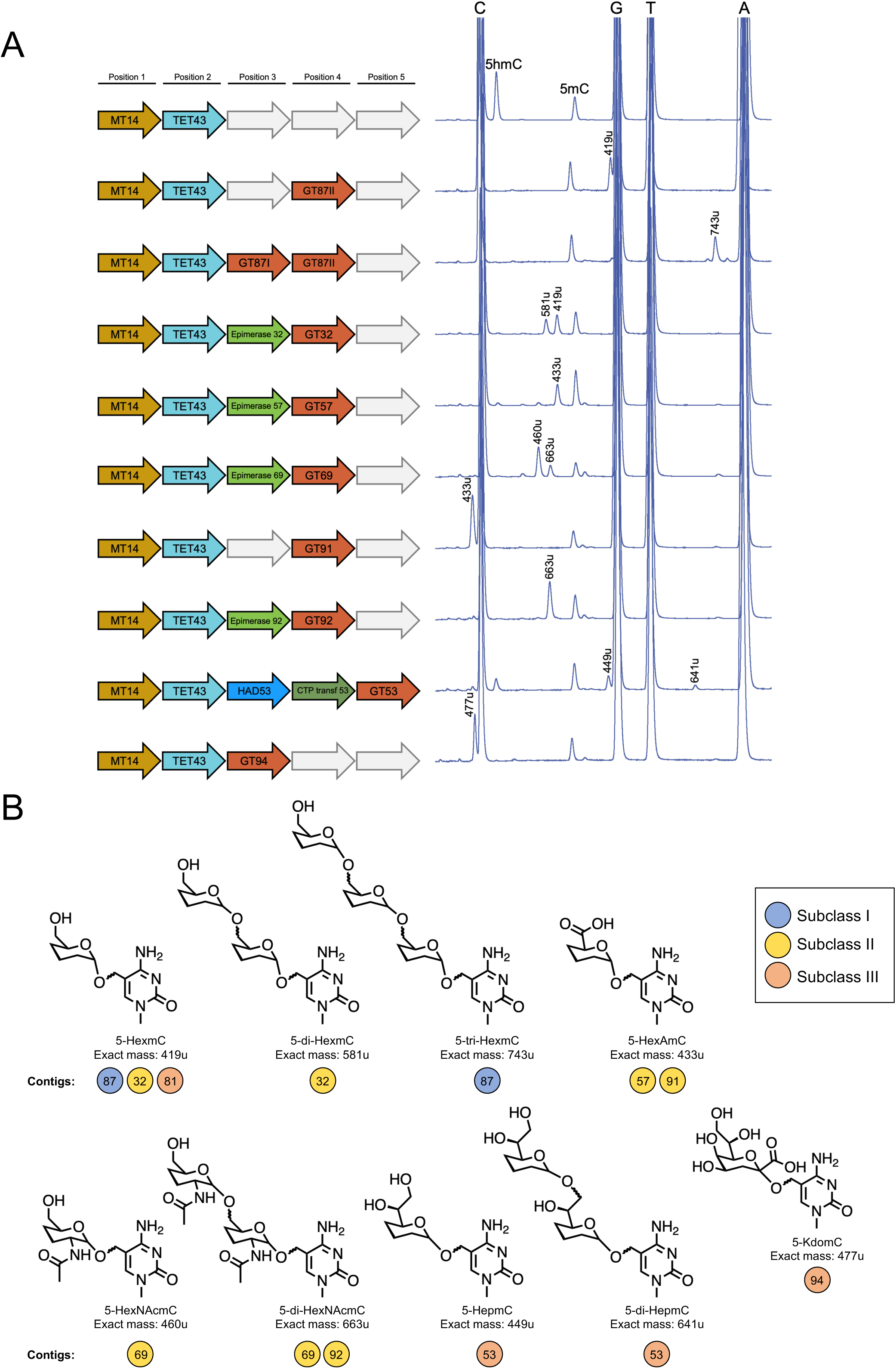
Novel cytosine sugar hypermodifications installed by phage DNA GTs. (A) UHPLC-MS analysis (right) of modified gDNA from E. coli expressing phage-encoded hypermodifying enzymes shown (left). Masses of compounds eluting within new LC peaks were used for identification of novel sugar modifications on 5mC. (B) Classes of new sugar products resulting from the biosynthetic activities of phage BGCs. Chemical structures of the sugar-modified nucleobase are shown. Colored circles represent contig number and subclass and indicate which sugar compounds were synthesized by each pathway.

For subclass I, we were able to successfully reconstitute the activity of the two adjacent GTs from pathway 87 (**Fig. S3A-B**). As previously observed from our work with the GTs from contigs 14 and 43 encoding a subclass I pathway, the downstream GT is responsible for installation of the first sugar moiety onto 5hmC whereas the upstream, 5mYOX-proximal GT in the pathway adds subsequent sugars to the growing polysaccharide hypermodification showing some level of synteny and functional conservation (18). In line with these initial observations, GTII alone from contig 87 produces a single hexose-5mC compound (419 amu) derived from 5hmC while expression of GTI alone did not result in any detectable modifications to 5mC and 5hmC (**Fig. S3C**). Co-expression of GTI and GTII produced a new hypermodified cytosine product trihexose-5mC (743 amu). The natural products from contig 87 enzymes provide further support for the position-specific roles of the two GTs in subclass I pathways. Moreover, these findings highlight the unique activity of GTI from contig 87 in successive installation of a mono- or disaccharide onto an existing monosaccharide-modified DNA base.

With only one exception (contig 91), we found that the enzymatic ability of GTs encoded in subclass II BGCs to install sugars onto 5hmC was strictly dependent on co-expression of their corresponding putative sugar epimerase (**Figs. 3A, S2 and S4**). As such, the favored hypothesis for subclass II GTs is that their activity depends on the availability of an appropriate activated sugar-nucleotide substrate molecule that is synthesized by their partner epimerase. Some subclass II-encoded GTs were also processive, producing both mono- and disaccharide modified 5mC. Two examples include the GTs from contigs 32 and 69, which are apparent processive hexose (Hex)- and *N*-acetylhexosamine (HexNAc)-transferases, respectively (**Figs. S2 and S4**). Co-expression of epimerase 32 and GT32 resulted in the biosynthesis of 5-hexosemethylcytosine (5-HexmC, 419 amu) and 5-di-HexmC (581 amu) from 5hmC precursor bases. By contrast, co-expression of epimerase 69 and GT69 produced 5-*N*-acetylhexosaminemethylcytosine (5-HexNAcmC, 460 amu) and 5-di-HexNAcmC (663 amu). A similar 5-di-HexNAcmC-containing product peak at the identical retention time was synthesized by epimerase and GT from contig 92, but the monosaccharide modified product was not observed (**Fig. 3A**). Contig 69 encodes an additional putative polyamine aminopropyl transferase (PAPT) enzyme that did not affect the accumulation of new hypermodification products in cells (**Fig. S2C**), suggesting that the substrate for this enzyme was not present in the *E. coli* metabolome. Contig 57 epimerase and GT co-expression and contig 91 GT alone both produced hexuronic acid (HexA)-modified cytosine (433 amu, **Fig. 3A**) eluting at different LC retention times, suggesting a possible epimerase-dependent isomeric difference between these two glycosylated nucleobase products. These findings support the hypothesis that most GTs encoded in subclass II BGCs coordinate with a partner epimerase enzyme, presumably to modify the stereochemistry of the pool of available sugar nucleotides in *E. coli* to provide the correct substrate for GT-mediated DNA hypermodification. However, GT activity may not be strictly dependent on epimerase association, as is the case for the GT from contig 91 (**Fig. 3A**). This may also be due, in part, to the bioavailability of preferred GT substrates within different host cells.

Of the subclass III GT-containing BGCs tested, only enzymes from contigs 53 and 94 yielded sugar-modified products (**Figs 3A** **and S5**). Our initial exploration into the GTs from contigs 36, 66, and 81 showed no significant activity, suggesting incomplete pathway reconstitution or unavailable substrates within our expression host, or a combination of both. Expression of GT94 alone was sufficient to produce a novel compound with a mass of 477 amu eluting at a relatively early LC retention time on reverse phase, supporting the prospective assignment of the sugar modification containing a carboxylate. Both observations were consistent with the assignment of a keto-deoxyoctonic (Kdo) sugar acid appendage (**Fig. 3A-B**). Contig 53 encodes a putative haloacid dehydrogenase (HAD)-like and aminoglycoside 3’-phosphotransferase (APH)-like bifunctional enzyme, cytidylyltransferase (CTP transf.), and GT-B fold-containing DNA GT with shared structural architecture with T4-βGT (**Figs. 1E** **and S5**). Expression of all three enzymes was essential for synthesis of two novel species corresponding to mono-heptose (Hep) (449u) and 5-di-HepmC (641 amu) (**Figs. 3A** **and S5**).

Collectively, the screening of twelve candidate phage BGCs revealed the biosynthesis of ten new cytosine sugar hypermodifications stemming from the *E. coli* metabolomic substrate pool.

### Phage GTs share structural features with eukaryotic and bacterial enzymes

Our discovery of diverse and complex glycans appended to cytosine raised questions about the origins of these newly identified phage GTs and their sugar substrates. Structural folding models of the tested C5-MT and 5mYOX-associated GTs were predicted using the AlphaFold (42) deep-learning network and revealed two general structural classes with GT-A and GT-B-like folds, in agreement with our phylogenetic analyses (**Figs. 1F, 4, and S1**). Using these predictive models, we probed the Protein Data Bank (PDB) and AlphaFold Database (AFDB) Proteome for structural homologs using Foldseek (43).

**Figure 4.**
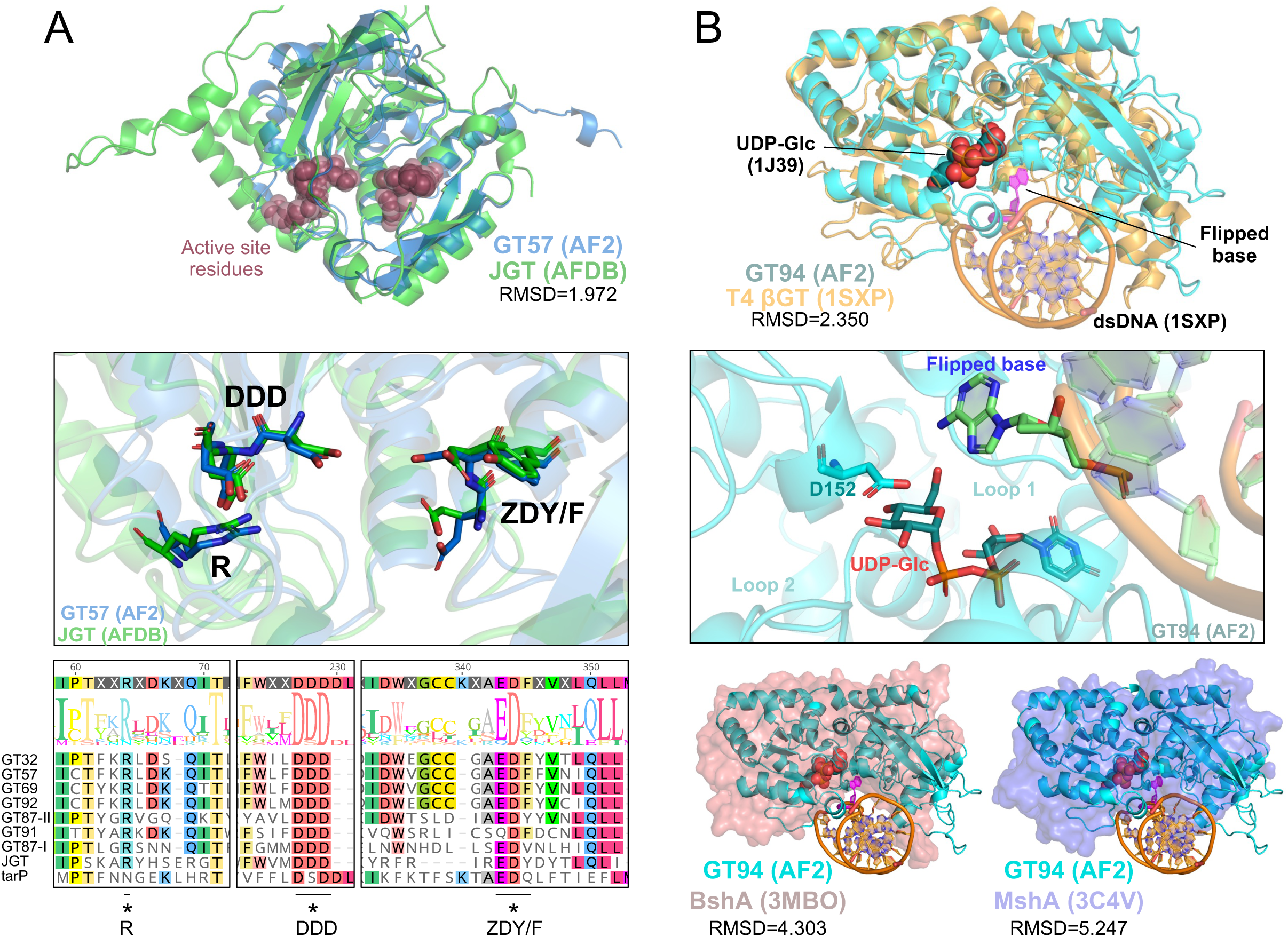
Predicted structural similarities between phage and cellular GTs. (A) Sequence and structural alignments of phage GT-A-like enzymes and their top cellular cellular homologs. Top: GT57 (light blue) as a representative of the GT-A-like enzymes is shown aligned with *T. cruzi* JGT (light green). Both predicted structures are illustrated in cartoon form with putative active site residues shown as red spheres. Middle: Magnified view of the aligned active site with key structural and catalytic residues highlighted as sticks. The R…DDD…EDY/F conserved GT-A active site motifs are well aligned between the two structures and labeled accordingly. Bottom: Multiple sequence alignment of all phage GT-A-like enzymes with top structural homologs from the PDB (tarP) and AFDB proteome (JGT) datasets. Aligned regions containing the conserved active site motifs highlighted above are indicated with asterisks. (B) Structural alignments of phage GT-B-like enzymes with top homologs from the PDB. Top: GT94 (cyan) as a representative of the GT-B hypermodifying enzymes is shown aligned with UDP-glucose (1J39) and DNA-bound (1SXP) structures of T4 βGT. UDP-glucose (UDP-Glc) is shown as red spheres while the position of the flipped mismatched G residue from the dsDNA duplex is highlighted in magenta. Middle: Magnified view of the putative GT94 (cyan) active site with the putative catalytic residue D152, UDP-Glc substrate, and flipped base shown in stick form. Two flexible loop regions flanking the glucose moiety of the activated sugar nucleotide are indicated. Bottom: Alignment of GT94 with additional top hits from the PDB, including BshA from *Bacillus anthracis* (3MBO) and MshA from *Corynebacterium glutamicum* (3C4V). Both bacterial GTs are shown as surface representations aligned with GT94 shown in cartoon form. UDP-Glc (red spheres) and dsDNA (orange helices with magenta flipped base) are modeled from 1J39 and 1SXP structures of T4 βGT, respectively to highlight overlapping predicted substrate binding regions for each enzyme.

Structure-guided searches of the AFDB using the phage GT-A structure predictions as query yielded consistent strong predicted matches (E-values = 10^-12^ to 10^-13^) with an uncharacterized protein from *Trypanosoma cruzi* (**Table S1**). Examination of proteins in UniProt (44) with 90% sequence identity to this hit revealed that this enzyme is a close homolog of the base J-associated glucosyltransferase (JGT) (45), providing further supporting evidence that the GT-A fold phage GTs directly transfer sugar moieties to DNA (**Fig. 4A**, **Table 2**). A MSA of the phage enzymes with the *T. cruzi* JGT revealed limited (∼9 % identity, **Fig. S6**) sequence similarity, suggesting an ancient structure-function relationship between the phage and eukaryotic DNA GTs. The same structure-guided search against experimental structural models in the Protein Data Bank (PDB100) returned top hits (E-value = 10^-5^ to 10^-6^) against tarP (**Fig. 4A, Tables 1 and S1**), a prophage-encoded GlcNAc-transferase for alternative biosynthesis of wall teichoic acid (WTA) in *Staphylococcus aureus* (46). TarP-mediated β-O-GlcNAcylation of ribitol-phosphate (RboP) subunits in WTA alters susceptibility of the bacterial cell to phage infection and modulates *S. aureus* immune evasion in mammalian hosts (46–48). Additionally, the chemistry of tarP-synthesized sugar conjugates matches those of the bacteriophage-encoded GTs – both transfer sugars to nucleophilic hydroxyl groups on RboP polymers or 5hmC acceptor substrates in DNA (46).

**Table 2.** New sugar hypermodifications installed by phage-encoded BGCs.

| Subclass | Contig # | Enzymes | Product 1 |  |  |  | Product 2 |  |  |  |
| --- | --- | --- | --- | --- | --- | --- | --- | --- | --- | --- |
|  |  |  | LC Retention Time | Mass (u) | Sugar(s) Added | Nucleoside Product | LC Retention Time | Mass (u) | Sugar Added | Nucleoside Product |
| I | 87 | GTII | 4.58 | 419 | Hexose | 5-HexmC |  |  |  |  |
| I | 87 | GTI+GTII | 6.77 | 743 | TriHexose | 5-tri-HexmC |  |  |  |  |
| II | 32 | Epimerase+GT | 3.4 | 419 | Hexose | 5-HexmC | 3.2 | 581 | DiHexose | 5-di-HexmC |
| II | 57 | Epimerase+GT | 3.46 | 433 | HexA | 5-HexAmC |  |  |  |  |
| II | 69 | Epimerase+GT | 3 | 460 | HexNAc | 5-HexNAcmC | 3.3 | 663 | DiHexNAc | 5-di-HexNAcmC |
| II | 91 | GT | 1.75 | 433 | HexA | 5-HexAmC |  |  |  |  |
| II | 92 | Epimerase+GT | 3.3 | 663 | DiHexNAc | 5-di-HexNAcmC |  |  |  |  |
| III | 53 | HAD+CTPtransf+GT | 4.54 | 449 | Heptose | 5-HepmC | 6.6 | 641 | DiHeptose | 5-di-Hep-mC |
| III | 94 | GT | 1.69 | 477 | Kdo | 5-KdomC |  |  |  |  |

Two notable exceptions among the phage-encoded GT-A fold enzymes were GT14I and GT43I from subclass I (**Fig. 1F**). Both are the first of two successive GTs which append secondary sugars to the growing glycan appended to 5hmC (18). Structural models of these two enzymes are nearly identical (RMSD= 0.342) and most closely resemble (E-values = 10^-7^ to 10^-8^) the predicted fold of an endogenous GT from *Enterococcus faecium* (general stress protein A, family 8 glycosyltransferase UniProt ID: A0A132Z517). This enzyme has a putative lipopolysaccharide (LPS) 1,3-galactosyltransferase domain and is related to the waaJ (RMSD=2.017) and waaO (RMSD=1.738) LPS glucosyl- and galactosyl-transferases, respectively, in *E. coli* (49–51). From the PDB, these phage enzymes had a top structural match with a major capsid protein glycotransferase from the giant algal virus PBCV-1 (**Tables 1 and S1**). Collectively, these structure-guided searches of the GT-A fold-containing phage enzymes underscore the ancient evolutionary exchange and functional repurposing of enzymes across prokaryotic and eukaryotic organisms and their viral pathogens.

We next performed the same searches using the GT-B fold enzymes, which are all GTs encoded in subclass III BGCs (*i.e.*, GT53, GT66, GT94 **Fig. 1E-F**). All three of the subclass III GTs shared some predicted structural folds with experimental structures of T4-βGT (E-values = 10^-5^ to 10^-7^, **Table S1**), in agreement with both the predicted GT-B fold and the ability of these enzymes to act on DNA substrates. The subclass III models position the predicted GT catalytic residues within a charged groove, which could function to bind dsDNA in a highly similar manner to T4-βGT (**Figs. 4B** **and S7**). Two loop regions unique to GT94 and GT53 flanked the predicted sugar binding pocket adjacent to the putative catalytic aspartate, possibly providing flexibility and increased solvent-exposed cavity space for accommodating more bulky sugar moieties (*e.g.,* Kdo, heptose) compared to glucose substrates preferred by T4-βGT (**Figs. 4B** **and S8**). In addition to T4-βGT, the subclass III GTs resembled structural models in the PDB corresponding to multiple bacterial GTs, including BshA (N-acetyl-α-D-glucosaminyl L-malate synthase), MshA (D-inositol 3-phosphate glycosyltransferase), from multiple species and PglH (GalNAc-α-(1,4)-GalNAc-α-(1, 3)-diNAcBac-PP-undecaprenol α-1,4-N-acetyl-D-galactosaminyltransferase) from *Campylobacter jejuni* (E-values = 10^-5^ to 10^-7^, **Fig. 4B, Tables 2 and S1**). Comparison with the top hits from AFDB Proteome revealed a pattern of structural similarities with bacterial and eukaryotic GT-B fold-containing enzymes that function primarily on glycolipid or small molecule precursors with free hydroxyl groups (**Tables 2 and S1**).

### Expanding the search for DNA sugar hypermodification pathways in the microbial metagenome

Using the GT-A and GT-B fold-containing phage enzymes (**Figs. 1, 4, and S1**) we generated hidden Markov model (HMM) profiles to re-probe metavirome and metagenome databases for related enzymes and their neighboring gene clusters. We expanded upon our initial search to include diverse microbial metagenomes by ecosystem curated within the Joint Genome Institute (JGI) Integrated Microbial Genomes and Microbiomes (IMG/M) (52), the gut phage database (GPD) (53), RNA viruses in the metatranscriptome (54), and all viral, archaeal, and enterobacterial reference genomes available in NCBI (**Table S2,** **Fig. 5A**). All hits from our GT HMM searches were retrieved with their surrounding gene neighborhoods, including ten predicted coding sequence regions upstream and downstream of the target gene (**Figs. 5** **and S9**). All coding regions in the retrieved contigs were annotated using HMMs from Pfam (55), TIGRFAM (56), REBASE (57), Carbohydrate Active Enzymes (CAZy) (58), and the prokaryotic antiviral defense locator (PADLOC) (59) and DefenseFinder (60) protein domain databases (**Fig. 5A**). This allowed us to make predictions about the roles of newly identified GTs in the context of their adjacent biosynthetic pathways.

**Figure 5.**
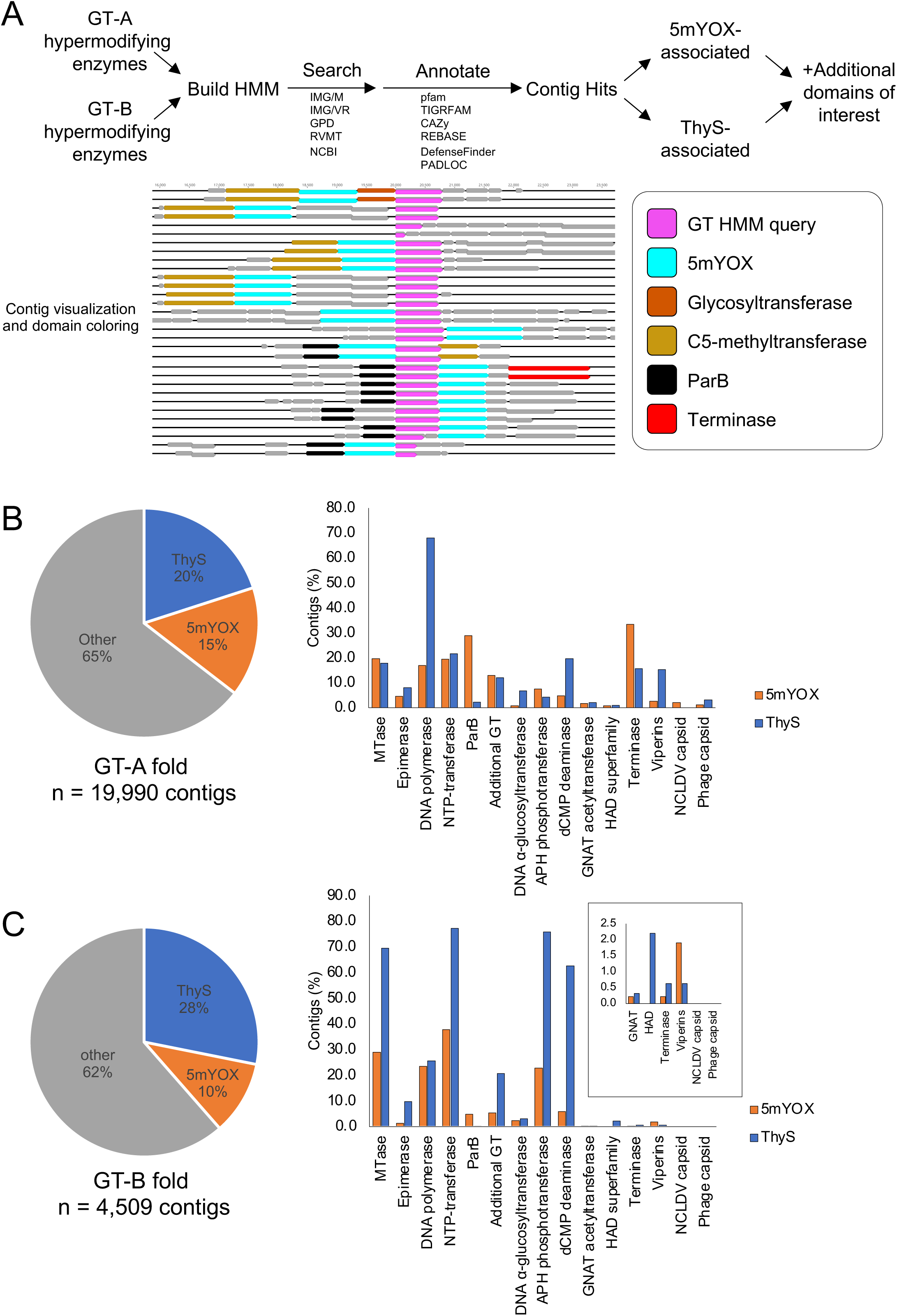
The expanding landscape of cytosine sugar hypermodification pathways in the metavirome. (A) Workflow for genomic and metagenomic database mining and discovery of new GT-containing pathways. (B) Summary of results from GT-A HMM mining. Left: pie chart showing the breakdown of ThyS and 5mYOX-associated contigs. Right: percentage of contigs containing GT-A hits and additional domain of interest, plotted by 5mYOX (orange) or ThyS (blue) association. (C) Summary of results from GT-B HMM mining. Left: pie chart showing the breakdown of ThyS and 5mYOX-associated contigs. Right: percentage of contigs containing GT-B hits and additional domain of interest, plotted by 5mYOX (orange) or ThyS (blue) association.

Our searches yielded approximately 25,000 new GT-containing contigs from all databases probed (**Figs. 5B-C** **and S9**). The vast majority of these contigs (∼95%) were identified within the latest release of the IMG/VR database confirming their viral origins (**Fig. S9**). The remaining ∼5% of assemblies were retrieved from IMG/M environment-specific metagenomes, the GPD, or from bacterial, archaeal, or phage genomes in NCBI. Most of the GT-A or GT-B-containing contigs (n=15,945, ∼65%) were not found neighboring another annotated base modification pathway enzyme (e.g., 5mYOX, ThyS) and are therefore less likely to be obviously involved in DNA hypermodification. Although we cannot rule out GTs coordinating with other base modification enzymes or 5mYOX/ThyS encoded further away in the phage genome, clustering of intact DNA hypermodification pathways seems to be a common feature in phage genomes (16, 18, 24). GT-A fold enzymes were more abundant in the metagenome (**Figs. 5** **and S9**, n=19,990 contigs, ∼82% of total), in agreement with the relative proportion of GTs in our initial limited dataset (18) (**Fig. 1**). Pathways containing a GT-B-like enzyme were more often associated with thymidylate synthase (ThyS, ∼28%) compared to 5mYOX (∼10%) reflecting a primary association with pre-replicative DNA hypermodification pathways (**Fig. 5C**). By contrast, the GT-A-like enzymes were more evenly distributed between 5mYOX and ThyS (**Fig. 5B**).

In addition to neighboring 5mYOX and ThyS enzymes, newly identified GTs were found adjacent to ORFs with annotated functions in DNA synthesis and modification (*e.g.,* DNA polymerase, DNA partitioning ParB-like proteins, helicases, methyltransferases), phage replication and DNA translocation proteins (*e.g.,* terminase, gp45, clamp) and additional sugar modifying enzymes (*e.g.,* epimerase, secondary GTs, APH phosphotransferase) (**Figs. 5B-C, S9, and S10**). For both GT-A and GT-B-associated pathways, only ∼20-30% of the 5mYOX-containing contigs also had a methyltransferase nearby (**Fig. 5B-C**), suggesting that many 5mYOXs are probably 5-hydroxymethyluracil (5hmU)-synthesizing rather than 5hmC. Only six contigs were identified that contained both a putative ThyS and 5mYOX enzyme (**Fig. S11**). In all cases, both annotated genes were within 10 kb of a GT-A HMM query hit, suggesting by association that there are relatively rare phage genomes that have the potential for carrying out both pre- and post-replicative installment of 5hmC/U hypermodification precursor bases. Certain gene annotations (*e.g.,* ParB, dCMP deaminase) were specifically associated with 5mYOX or ThyS-containing pathways regardless of GT fold present (**Fig. 5B-C, S10**). Intriguingly, many of the GT- and 5mYOX/ThyS-containing pathways harbored genes with predicted functions in prokaryotic immune systems, including annotated viperins (pVip), abortive infection (Abi) systems, components of the Gabija (Gaj) nucleotide sensing defense system, and many restriction nucleases (**Figs. 5B-C, S10**). A small number of pathways (n=65) from IMG/VR were found containing a GT-A-like enzyme and a putative 5mYOX and a nucleo-cytoplasmic large DNA virus (NCLDV) capsid protein (61), suggesting the presence of related DNA hypermodification pathways in the genomes of giant eukaryotic viruses (**Figs. 5B** **and S12**).

These observations that i) many microbial metagenome search hits were found in gene neighborhoods that do not contain pathways that are obviously associated with DNA hypermodification (**Fig. 5B-C**) and ii) GT structural homologs (**Fig. 4**) and similar GT sequences from cellular organisms have been identified in all three domains of life (**Table S3, Fig. S9**) further support the hypothesis that the phage-encoded GTs have cellular homologs that have been repurposed throughout evolution to glycosylate diverse substrate acceptor molecules. Moreover, the functional and structural overlaps between phage GTs and those of endogenous eukaryotic and bacterial organisms suggest an evolutionary adaptation that has been shaped by the dynamic interplay between phage genome defense and bacterial immune systems.

## DISCUSSION

The primary finding from our study is that diverse sugar structures are installed at the 5-carbon position of cytosine by enzymes encoded from new DNA hypermodification pathways in viral genomes. These enzymes appear to be structurally related to those present in bacterial and eukaryotic cells and have been repurposed throughout host-virus evolution. These enigmatic DNA glycans expand our understanding of the plasticity and chemical tolerance of nucleic acids and underscore the vast unexplored biology of organisms that have evolved to use unconventional modifications for genomic regulatory functions (62). To discover and characterize new compounds, we developed and optimized a HT biosynthetic pathway reassembly platform for co-expression of metavirome enzymes and isolation of their hypermodified DNA products. As our analytical biochemistry and bioinformatic approaches converged, we revealed potential for evolutionary connections between the newly identified phage DNA hypermodification pathways and endogenous lipopolysaccharide, wall teichoic acid, glycolipid and small molecule biosynthesis pathways of bacterial host organisms. These biological similarities allowed us to expand our search for new biosynthetic gene clusters in the phage metavirome and develop hypotheses about the roles of sugar-coated DNA in the phage infection cycle.

### Cell-based BGC assembly for the discovery of novel nucleic acid modifications

To assess the activities and products of the enzymes in our BGCs, we sought to develop a scalable and unbiased platform to provide these enzymes with a pool of potential substrates and a suitable acceptor molecule. We knew from our previous co-expression methodology that phage-encoded C5-MT, 5mYOX, and GTs could act directly on *E. coli* gDNA (18) and thus aimed to reassemble entire core components of the discovered BGCs in a single expression vector. Our approach allows for semi-automated HT assembly of BGCs in any combination without concern for ineffective recognition of native promoter sequences. Type II-S RE-based assembly approaches have been successfully developed to reassemble functional BGCs in *E. coli* (63, 64) and other organisms (65–68) for natural product biosynthesis and pathway analyses. The system developed here offers a specific advantage for investigation of novel predicted nucleic acid-modifying pathways, as the non-nucleated *E. coli* genomic DNA offers a readily available template for enzymes operating in diverse sequence contexts. Moreover, the downstream steps (*i.e.,* whole-plasmid sequencing, gDNA isolation, on-bead clean-up and digestion to free nucleosides, UHPLC-MS analysis) are coordinated in 96-well plate format for HT assessment of pathway functions and nucleic acid hypermodification discovery. With minimal adaptation, we anticipate this platform being amenable for pathways predicted to install novel RNA modifications. Moreover, our platform design allows for customized approaches to probe pathways for small molecule biosynthesis, protein or lipid modifications.

### Origins of phage DNA hypermodifying enzymes

Viruses have undoubtedly shaped the evolution of all cellular life (69). Recently, Krupovic, Dolja, and Koonin predicted that the virome of the last eukaryotic common ancestor evolved, in part, from bacterial viruses (70). Our identification of structural homologs of the phage DNA GTs from bacterial and eukaryotic species (**Figs. 4, S6, and** **S9; Tables 1 and S1**) is supported by the apparently ancient fold of GTs in evolutionary space (71, 72). The “uncharacterized protein” (UniProt Q4E4F0, putative JGT) from *T. cruzi* as a top hit for the GT-A phage GTs suggests that divergent eukaryotic DNA GT homologs are present in the biosphere. Indeed, a blastp search using the putative base J glucosyltransferase from *T. cruzi* revealed many homologs primarily belonging to members of the *Trypanosoma*, *Leishmania*, and several other trypanosome genera, but also cyanobacteria (*Spirulina* sp.) and predatory bacteria (*Bdellovibrio* sp.) (**Table S3**). These findings highlight the utility of structure-guided searches to investigate intra-domain enzymes with related folds that share limited sequence similarity. Moreover, the structural similarities with a prophage-encoded GT (TarP) hints at the possibility that these post-replicative DNA hypermodification enzymes have structural homologs encoded within mobile genetic elements, which supports the hypothesis of virus-mediated gene transfer between, or acquisition from, evolutionarily diverse organisms. It is also possible that phage-encoded DNA-hypermodifying GTs evolved from prophage-like elements with primary functions in superinfection exclusion via remodeling of the bacterial cell surface (46, 48). Evidence for phage acquisition and repurposing of host defense system genes for pro-viral functions has been recently reported (73) and supports this potential mechanism of GT origins and evolution. The discovery of putative anti-phage defense system gene annotations in the newly identified GT-associated pathways (**Figs. 5** **and S10**) is consistent with the presence of defense systems (*e.g.,* PARIS, viral Cas homologs) in the genomes of phage and phage satellites, and the transfer of this pathway information to host cells via prophage integration (74). Antiviral elements encoded in viral satellites and integrated into host genomes are also apparently abundant in the genomes of unicellular eukaryotic organisms (75, 76). The continued discovery, functional characterization, and evolutionary reconstruction of DNA GTs will further clarify the origins and biological roles of these unique enzymes.

Our findings further demonstrate that the activity and substrate selectivity of a candidate phage GT are, in part, determined by the contig and pathway architecture and interdependence on neighboring enzymes (**Figs. 3** **and S2-S5**). This neighborhood interdependence further supports the hypothesis that intact BGCs could be preferentially inherited vertically or shuttled between virus and host, preserving the observed synteny across different GT-containing subclasses (**Fig. 1C-E**). In this sense, the post-replicative hypermodification pathway cassette (**Fig. 1B**) could function as a mobile element acquired by phage during integration events and conferring adaptations to host range, new host defense systems encountered, regulation of gene expression, or other essential infection cycle functions (75, 76).

### Role of sugar-modified bases during infection

Sugar hypermodifications on phage DNA can block endonucleolytic cleavage by host restriction enzymes (15, 25, 26, 31, 34). However, these sugars are typically installed at high sequence coverage across the phage genome resulting from pre-replicative ThyS-based modulation at the level of the cellular nucleotide pool (16, 25). By contrast, post-replicative 5mYOX-based sugar hypermodifications are determined by the sequence context of phage-encoded C5-MT enzymes (**Fig. 1B**). Most 5mYOX-coupled phage MTs have a GpC sequence motif preference (18), limiting the site-specific installation of sugars onto 5hmC by downstream GTs. It therefore seems unlikely that this post-replicative strategy would broadly provide effective genome-wide protection from host endonucleases targeting cytosine in other sequence contexts. We cannot exclude the possibility that additional motif specificity is provided by the GT binding of the DNA polymer, or that glycans shield the phage DNA from GpC-specific endonuclease activity. Moreover, we do not know whether 5mYOX-associated hypermodification enzymes are preferentially acting on phage DNA, the host genome, or both. With our cell-based pathway reassembly platform, we observe that moderate amounts of complex sugar hypermodifications are tolerated by *E. coli* when installed on the bacterial chromosome (**Figs. 3** **and S2-S5**). Sequencing of phage and host genomes during infection and base-resolution mapping of installed modifications will be necessary to reveal the distribution of sugars and position relative to functional genomic regions.

Many of the GTs from all subclasses share structural homology with bacterial GTs, specifically those involved in biosynthesis of cell surface molecules (*i.e.*, LPS, WTA) and glycolipids (**Fig. 4** **and Tables 1 and S1**). Kdo, Hep, and additional LPS core sugars are receptors for phage infection (77–81) and multiple host enzymes in LPS biosynthesis were identified in screens for phage receptors (82). Sequestration of LPS sugars through conjugation to DNA bases by phage-encoded GTs could indirectly affect receptor maturation or abundance at the host cell surface or serve to redirect sugar binding proteins from the host cell. Thus, phage or host DNA used as a template by the phage-encoded GTs would effectively act as a sugar and/or sugar-binding protein sponge during infection. Since Hep and Kdo are specialized and essential sugars for LPS maturation (83), sequestration of the activated sugar-nucleotide substrates could suppress receptor presence at the cell surface preventing attachment by extracellular phage in the surrounding environment as a new mechanism of superinfection exclusion (84). It is also possible that, in addition to their role in DNA hypermodification, the LPS pathway-related phage GTs retain contacts with membrane structures or with membrane-associated LPS precursors. Directing phage DNA to membranous compartments within the cell could facilitate genome replication (85, 86) or other critical stages of the infection cycle, as observed recently by Armbruster and colleagues (87). It is also possible that the host sugar metabolites are concentrated at the point of viral DNA replication and therefore used opportunistically in DNA hypermodification systems. How and whether changes in the abundances of host metabolites alter the activities of the phage GTs remain to be explored. The biophysical properties of site-specific complex DNA glycans and regulatory effects on gene expression patterns, recruitment of reader or eraser proteins, and restructuring of the bacterial nucleoid or viral genome for packaging remain to be investigated.

Interestingly, our GT HMM profile searches only yielded a small number of hits from sequenced phage isolates in NCBI that represent only a limited subset of tailed phage (**Fig. S13 and Table S4**). The only post-replicative pathways (*i.e.,* 5mYOX-associated) in this subset originated from uncultured *Caudovirales* metavirome sequences, whereas the remainder were exclusively ThyS-associated and found in cultured phage genomes. Whether pre- or post-replicative GT-containing hypermodification pathways are mutually exclusive in viral genomes remains an open question, though one could envision scenarios where encoding both may provide context-specific advantages to the phage during infection. Alternatively, prophage-encoded GTs could provide anti-phage activity by hypermodifying invading viral genomes with complex sugars that disrupt replication by interfering with viral polymerase elongation. Whether such bulky modifications – either pro- or anti-viral – prevent conventional sequencing of phage genomes and therefore identification of these pathways in the metavirome poses an evolving question and challenge for the study of unconventional DNA chemistry. Indeed, modified bases in phage genomes have obstructed sequencing efforts using short-read-based approaches, either due to polymerase stalling or structural changes to the DNA (88, 89).

The ability to distinguish self from non-self is one of the greatest challenges to both virus and host during infection. While many eukaryotic viruses will modify existing subcellular compartments to sequester their genomes during replication (90), phage have adopted unique strategies to separate their nucleic acids from the bacterial chromosome. One example is the nucleus-like compartment formed by jumbo phages to exclude antagonistic host factors from their replication compartments (91). Whether decoration with diverse sugars has a function in phage DNA condensation for partitioning into replication factories or for packaging into newly formed capsids remains to be investigated. Cytosine modifications at the C5 position can have a marked effect on the helical conformation and biophysical features of the larger DNA molecule (92). Future biophysical and structural studies will reveal the impact of bulky polar glycans on the architecture and physicochemical properties of hypermodified DNA.

The presence of new DNA hypermodifications resulting from phage encoded post-replicative BGCs expands our understanding of these pathways and the chemical possibilities tolerated by nucleic acids. The abundance of related enzymes and pathways in the metagenome suggests that DNA glycan hypermodifications may be more common than previously appreciated and play largely undefined roles in the biology of cellular and parasitic organisms, alike.

## MATERIALS AND METHODS

All enzymes, buffers, plasmids, and strains were obtained from New England Biolabs (NEB, Ipswich, MA) unless otherwise noted. Sequences of the native contigs, codon-optimized gene fragments, and destination vectors are included with the SI. PCR setup, amplicon normalization, and Golden Gate Assembly (GGA) reaction setup steps were performed using custom python scripts operated on an OT-2 (Opentrons, Brooklyn, NY). Python packages used for data analysis include SciPy, Pandas, Seaborn, and Matplotlib.

### Design of BGC destination expression vectors pSL003 and pSL126

Entry vectors pSL001 and pSL126 for the *E. coli*-based expression platform were generated using NEBuilder® HiFi DNA assembly using pET28a (Millipore Sigma, Burlington, MA) and pGGA as templates. Our initial design consisted of a pBR322 plasmid backbone containing a KanR marker and lacI repressor. We built the backbone with a lacZ cassette that would be released upon treatment with BsaI for use in GGA. Inspired by the Biopart Assembly Standard for Idempotent Cloning (BASIC) method (93), we designed a process for rapid operon reconstitution that appended construction functionality in a single step. We designed primers to amplify our parts using Q5-High Fidelity Polymerase that would afford our desired overhangs for assembly in a GGA reaction mix. In addition, the primers appended barcodes that could be used as annealing sites for diagnostic amplifications and sequencing. In addition, we moved from a pMB1 to a p15A origin of replication to reduce plasmid copy number, in an effort to increase BGC stability and reduce looping out of the repetitive T7 promoter-terminator regions which we observed during system development and validation. For the combinatorial BGC cell-based assays, we developed a plasmid backbone containing MT14 and TET43 flanking the f1 origin of replication and KanR genes, such that in the event of looping out the plasmid would be deficient in critical elements necessary for stability and propagation.

### Cloning of individual genes from initial GT-containing pathways

Full-length genes from candidate BGCs were codon optimized for expression in *E. coli* and domesticated to remove any BsaI restriction enzyme cut sites. Gene fragments were chemically synthesized (Twist Bioscience, South San Francisco, CA) with overhangs for Golden Gate Assembly into the pSL003 destination vector. GGA was performed using 75 ng of each gene fragment insert and 10 ng of plasmid under standard reaction conditions according to the NEB BsaI-HF®v2 protocol for single gene assembly (full protocol available at: www.neb.com/en-us/protocols/2018/10/02/golden-gate-assembly-protocol-for-using-neb-golden-gate-assembly-mix-e1601). GGA were conducted using 760 U T4 DNA ligase, 22.5 U BsaI-HF®v2, and 1 X T4 DNA Ligase Reaction Buffer (50 mM Tris-HCl, 10 mM MgCl2, 1 mM ATP, 10 mM DTT at pH 7.5). Assembled vectors were transformed into NEB® 5-alpha competent *E. coli* cells (C2987) and verified by Sanger sequencing and colony PCR for correct amplicon size (**Fig. S14**).

### BGC assembly for pathway expression in *E. coli*

Individual genes for multiple piece assemblies were first PCR amplified in 50 µL reactions using Q5® Hot Start High-Fidelity 2 X Master Mix using 0.5 µM of each primer. Primers annealed to conserved regions of the individual gene vectors and appended barcodes and sequences enabling high fidelity assembly. In the leave-one-out tests of neighboring gene dependency, the empty regions for omitted genes were replaced with pET28a multiple cloning site inserts. An OT-2 liquid handling robotics instrument (Opentrons, Brooklyn, NY) was used to assemble template and appropriate primers for each reaction. PCRs were performed using an initial denaturation at 98°C for 30 seconds followed by 30 cycles of 98°C for 5 seconds, 60°C for 10 seconds, and 72°C for 2 minutes and a final extension step at 72°C for 2 minutes. Amplicons were purified using 1.8X volume of SPRIselect (Beckman Coulter, Brea, CA) and a KingFisher Flex (ThermoFisher Scientific, Waltham, MA). Beads were washed twice with 300 µL 80 % ethanol and eluted in 50 µL water. Amplicon quality was assessed using Latitude™ HT Precast, 1.2% SeaKem® LE Plus Agarose gel (Lonza, Basel, CH). Recovered amplicon abundance was measured in 384-well plates using a SpectraMax Microplate Reader (Molecular Devices, San Jose, CA) with a Quant-iT™ Broad-Range dsDNA Assay Kit (ThermoFisher Scientific, Waltham, MA). All amplicons were diluted to 30 nM using customs scripts on the OT-2 instrument. Final BGC assembly reactions were performed using 10 ng of destination vector (pSL003 or pSL126) and 2.2 nM of each amplified insert following the NEB BsaI-HF®v2 protocol for multiple gene assembly. Individual amplicons were consolidated for assembly using an OT-2 based on BGC design. Reactions were cycled between 37°C and 16°C, each at 5 minutes, for a total of 35 cycles followed by a 5-minute incubation at 55 °C and cooling to 4°C. Assembled expression vectors were transformed into T7 Express Competent *E. coli* (NEB, C2566) and transformed cells were plated on selective Kan LB agar plates and incubated overnight at 30 °C.

### Expression of BGC assemblies in *E. coli*

Four to six colonies per design were transferred into deep-well plates containing 250 µL LB Kan. Plates were incubated with shaking overnight at 30 °C. The following day, cultures were diluted with 250 µL LB and 10 µL was used to inoculate a fresh plate containing 500 µL LB Kan in each well. Plates were incubated with shaking for five hours at 30°C before IPTG was added to 400 µM by being delivered with 25 µL LB. The plates were incubated at 18 °C overnight. The following day the cultures were transferred to KingFisher deep well plates (ThermoFisher), spun down at 2500 g for 10 min, and media removed by inversion. The same colonies used for starting the liquid expression cultures were used for cPCR verification of insert position and size (**Fig. S14**). PCR amplicons were separated and visualize using agarose gel electrophoresis and staining with ethidium bromide to verify product size and homogeneity.

### Purification of modified *E. coli* gDNA

Pellets were then resuspended with 95 µL PBS and 5 µL Proteinase K using a Thermomixer. Once achieving homogeneity, 95 µL Tissue Lysis Buffer from Monarch® Genomic DNA Purification Kit and 5 µL Monarch® RNase A was added and the plate was incubated with shaking at 1200 rpm on a Thermomixer at 56 °C for 4 hours until the lysates had clarified. 50 µL of SPRIselect and 100 µL PEG binding buffer (2.5 M NaCl, 20 % PEG 8000 w/v, 10 mM Tris pH 8.0) was added and the mixture was gently shaken at 800 rpm on a Thermomixer for 10 minutes. gDNA was then purified using a KingFisher Flex through three 300 µL 80 % ethanol washes followed by elution in 97 µL water containing 3 µL Monarch® RNase A leaving the SPRI beads in the elution. The elution plate was left on the benchtop overnight to digest the contaminating RNA. The following day, 100 µL PEG binding buffer was added and the bound gDNA was washed twice with 300 µL 80 % ethanol and eluted into 100 µL water with the beads being left in the elution.

### Enzymatic digestion of *E. coli* gDNA to free nucleosides

The eluted gDNA was resuspended by pipetting to make the beads and elution as homogenous as possible before 20 µL was used in a 40 µL digest using 1 µL of nucleoside digestion mix and 4 µL 10X buffer following the protocol for Nucleoside Digestion Mix (NEB) in an unskirted PCR plate. The mixture was incubated at 37 °C overnight on a thermocycler, filtered using AcroPrep Advance 96 Well 0.2 µm Supor Short Tip Natural PP (Pall Corporation, Port Washington, NY) into a skirted PCR plate. The plate was then covered with a Zone-Free™ Sealing Film (Excel Scientific, Victorville, CA) for UHPLC analysis. Samples for sugar mass analysis were generated by digesting 1.5 µg isolated gDNA in a 30 µL Nucleoside Digest reaction with 1 µL enzyme followed by filtration and 15 µL injected on an ultra-high performance liquid chromatography (UHPLC) instrument equipped with a mass spectrometer (MS).

### *In vitro* labeling of T4 *gt^-/-^* gDNA

T4 *gt^-/-^* gDNA containing all 5hmC and background levels of 5GlcαmC (kindly provided by Peter Weigele) was used for *in vitro* enzymatic labeling with recombinant T4 βGT for the UHPLC validation experiments shown in **Fig. 2B**. T4 *gt^-/-^* mutant phage was produced and isolated as described previously (94). Prior to *in vitro* glucosylation, phage gDNA was sheared to approximately 1500 base-pair fragments in 10 mM Tris pH 8.0 with 1 mM EDTA using a COVARIS S2 ultrasonicator instrument with the following settings: 5 % duty cycle, intensity 3 and 200 cycles/burst, as described previously (18). 1 µg of sheared gDNA was labeled in a 20 µL reaction with 10U recombinant T4 βGT in 1X NEB CutSmart buffer (50 mM potassium acetate, 20 mM Tris-acetate pH 7.9, 10 mM Mg(C₂H₃O₂)₂, 100 µg/ml BSA) supplemented with 2 mM UDP-Glc. Reactions were incubated at 37°C overnight followed by gDNA purification using the NEB Monarch® PCR and DNA Cleanup Kit. Total recovered gDNA was digested and injected for UHPLC as described above.

### UHPLC and LC-MS analysis of nucleosides

UHPLC analysis was performed on an Agilent 1290 Infinity II (Agilent, Santa Clara, CA) equipped with a G4212A diode array detector. UHPLC was carried out using a Waters XSelect HSS T3 XP column (2.1 × 100 mm, 2.5 µm) (Waters, Milford, MA) with buffer A being comprised of 10 mM ammonium acetate pH 4.5 and buffer B as methanol at a 0.6 mL/min constant flow rate and a 2 µL injection. The gradient was as follows: 1 % B for 1.5 min, 1-10 % B from 1.5 to 7.5 min, 90-100 % B from 7.5 to 9 min, hold at 100 % from 9 to 11 min, and 100-1 % B from 11 to 12 min before a 1.5 min hold prior to the next injection. LC-MS analysis of unknown peaks was performed using a similar setup, but with a 6120 single quadrupole mass detector in both positive and negative modes. Quantification was performed by injecting different amounts of digested DNA from phage Lambda (canonical DNA bases), phage Xp12 (C replaced with 5mC), T4 phage *gt^-/-^* C replaced with 5hmC), and T4-βGT treated T4 phage *gt^-/-^* (C replaced with 5-GlcβmC). Phage sequences were used to determine GC content (therefore A:G, G:C, C:A, etc) and scalars were experimentally determined to relatively quantify nucleosides based on absorbance at 260 nm. 5-GlcβmC was quantified based on converting 100 % of 5hmC using T4-βGT and UDP-glucose with T4 phage *gt^-/-^* and comparing the response factors (extinction coefficient multiplied by concentration) between substrate and product. Scalars calculated to normalize areas of various DNA species at 260 nm to result in areas that could be directly compared were as follows: A = 1; thymine = 1.65916; G = 1.22133; C = 2.05721; 5mC = 3.01239; 5hmC = 2.71319; and 5-GlcβmC = 2.828.

### Bioinformatic analyses

Amino acid sequence alignments for the GTs were generated in Geneious Prime (Dotmatics, Boston, MA) with default parameters for Multiple Alignments using Fast Fourier Transform (MAFFT) with a BLOSUM62 scoring matrix. The phylogenetic tree in Fig. 1F was generated using a MAFFT sequence alignment for phage-encoded GTs with FastTree (95) default parameters in Geneious Prime.

### Initial pathway annotations and synteny predictions

The initial set of GT-containing contigs was curated from our previously published dataset of C5-MT and 5mYOX-containing pathways (18). Conserved pathway architecture (Fig. 1) was predicted using the clinker web tool (cagecat.bioinformatics.nl/tools/clinker) with default parameters (96). Gene annotations were predicted using a combination of comparisons against pfam HMM profiles (55), HHpred (97), and Foldseek searches using AlphaFold2 structural models (42, 43).

### Structural model prediction and structure-guided homolog searches

Structural models were predicted using AlphaFold2 via ColabFold (98) with three recycles and five output models. The rank 1 model was used as input for Foldseek searches of the AlphaFold Proteome and PDB100 databases in 3Di/AA mode (43). Structural models and alignments of top Foldseek hits were visualized using PyMOL (Schrödinger, New York, NY). The top ten structural matches for each search are described in Table S1.

### GT HMM profile generation and re-probing of metagenomic databases

HMM profiles for GT-A and GT-B enzymes were assembled using HMMER3 (99) from MSAs generated in Geneious Prime (Dotmatics, Boston, MA). Individual HMMs were used as query for probing multiple metagenomic and reference sequence databases (**Table S2**) with an e-value cutoff of 10^-4^ for query HMM matches. Sequence contigs containing query matches were de-duplicated based on clustering by 100% sequence identity and clustered by sequence similarity to other hits. Sequences were retrieved with ten protein coding regions upstream and downstream of the query hit to reflect the genomic context and neighboring genes in the potential BGC. All protein coding regions in the retrieved contigs were annotated using HMMs from Pfam (55), TIGRFAM (56), REBASE (57), CAZy (58), DefenseFinder (60) and PADLOC (59) protein domain databases (**Fig. 5A**). Contigs containing protein coding region with additional domains of interest were extracted in Geneious using the parameters outlined in Table S5.

## ACKNOWLEDGEMENTS

The authors would like to thank members of the NEB Research community for thoughtful discussions and advice throughout this project, particularly past and current members of the Weigele, Correa, Taron, and Eaglesham labs. We would also like to thank the NEB sequencing core for assistance with Sanger sequencing and plasmid assembly verification, and Nan Dai for assistance with LC-MS sample processing. This work was supported by funding from New England Biolabs.

## SUPPLEMENTAL TABLE LEGENDS

**Table S1.** Expanded Foldseek results for phage-encoded hypermodification enzymes. AlphaFold structural models for each phage enzyme were used as query for probing protein structure databases using Foldseek, as described in the Materials and Methods. The top ten structural homologs from the PDB and AFDB Proteome databases are shown for each hypermodification enzyme. Expansion of the data shown in **Table 1**.

**Table S2.** Descriptions of the genomic and metagenomic databases searched in this study.

**Table S3.** *T. cruzi* base JGT blastp search results. The putative base J glucosyltransferase (JGT) from Trypanosoma cruzi strain CL Brener (UniProt Accession: Q4E4F0) was used as query for blastp search of all proteins in NCBI (blast.ncbi.nlm.nih.gov/Blast.cgi).

**Table S4.** List of viral genomes in NCBI with encoded GT-A structural homologs. Complete list including the selected representative genomes shown in **Fig. S10**.

**Table S5.** Gene annotation terms used to identify contigs with co-occurring domains of interest using Geneious Prime software.

---

**Figure S1.**
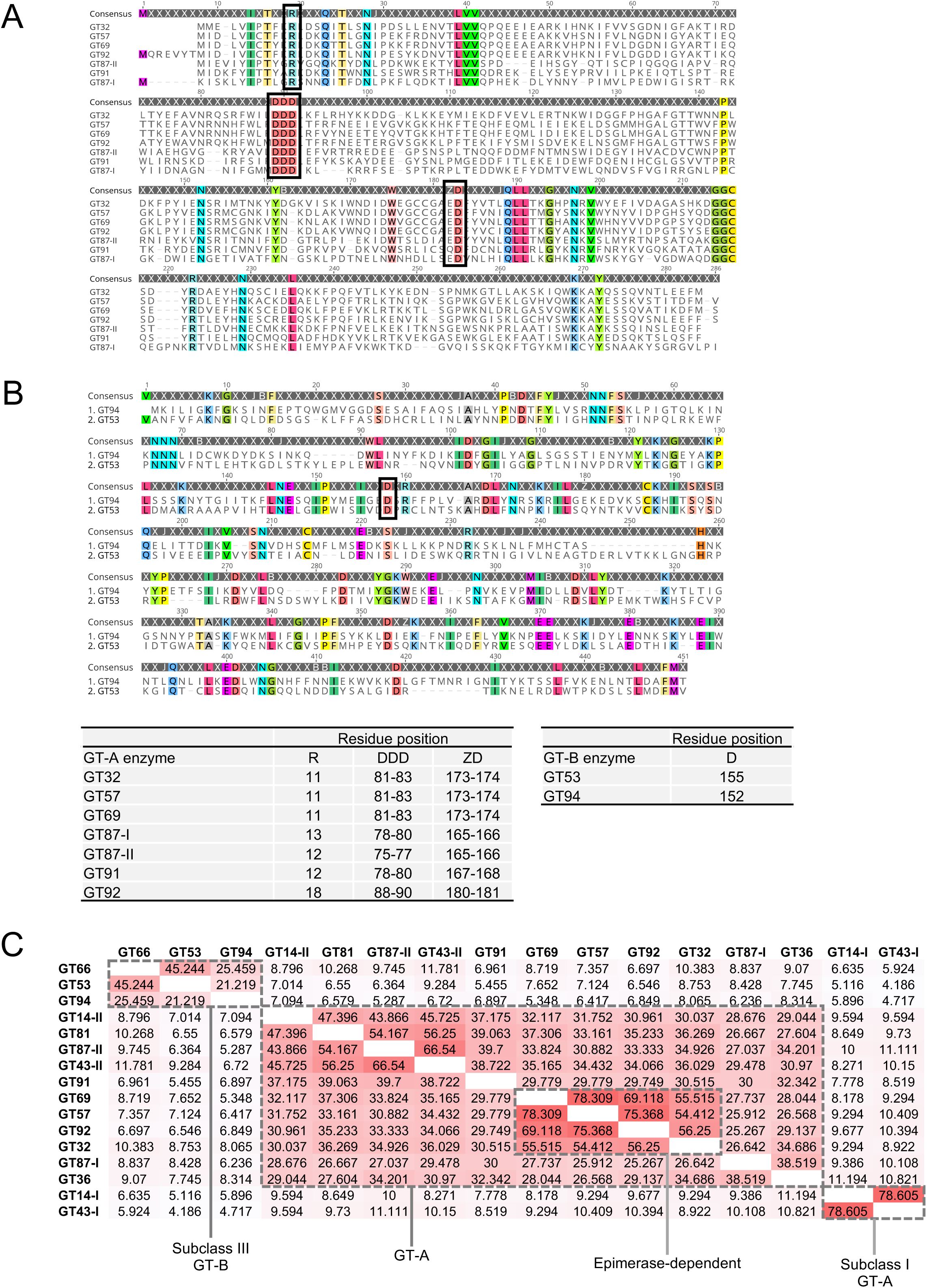
Multiple sequence alignment of 5mYOX-associated phage GTs with GT-A (A) and GT-B (B) folds. Multiple sequence alignments were generated as described in Materials and Methods. Conserved active site residues for the GT-A family is indicated with black boxes in A (24, 100). The proposed catalytic aspartate residue for the GT-B phage GTs is indicated with a black box in B, based on structural alignments with T4 BGT and overlap with the D100 catalytic residue (101) (see **Fig. 4B**). (C) Pairwise analysis of amino acid sequence similarity for all GTs tested in this study. Boxes are colored by increasing sequence similarity. Clusters of related GTs with functional significance are indicated with grey boxes.

**Figure S2.**
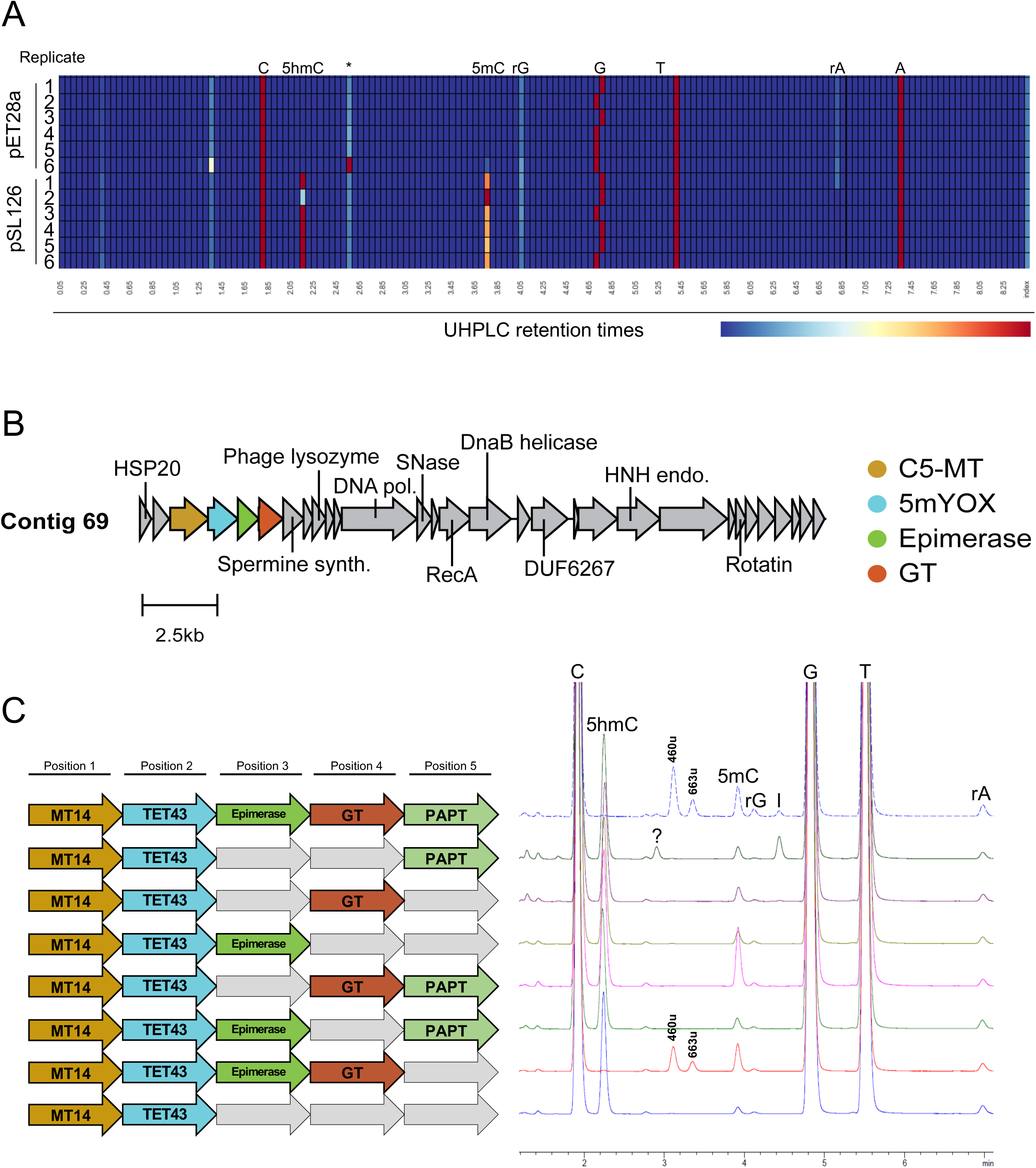
BGC assembly system validation using contig 69 enzymes as a test case. (A) Heat map of UHPLC results from digested gDNA samples fromT7 Express *E. coli* harboring empty vector controls (pET28a) or co-expressing MT14 and TET43 (pSL126) for synthesis of 5hmC hypermodification precursor residues. UHPLC retention times are shown below the heat map. Colored boxes represent peaks from the UHPLC traces. Heat map scale by color intensity is shown below and corresponds to total normalized area for peaks across retention time intervals. Results from six biological replicates are shown for each condition. The elution of a non-specific product peak of unknown mass is indicated with an asterisk. (B) Schematic of the contig 69 BGC. (C) Systematic analysis of contig 69 enzyme DNA hypermodifying activities in *E. coli*. Left: Different assemblies tested. Empty grey arrows represent positions in the pSL126 vector assembled with non-coding pET28a control amplicons to control for enzyme position and expression plasmid size. Right: UHPLC traces from digested *E. coli* gDNA nucleoside composition. Canonical based are indicated. Masses of new products synthetized in an epimerase at GT-dependent manner are shown above corresponding peaks. Elution peaks for ribonucleosides rG, rA, and inosine (I) are also indicated.

**Figure S3.**
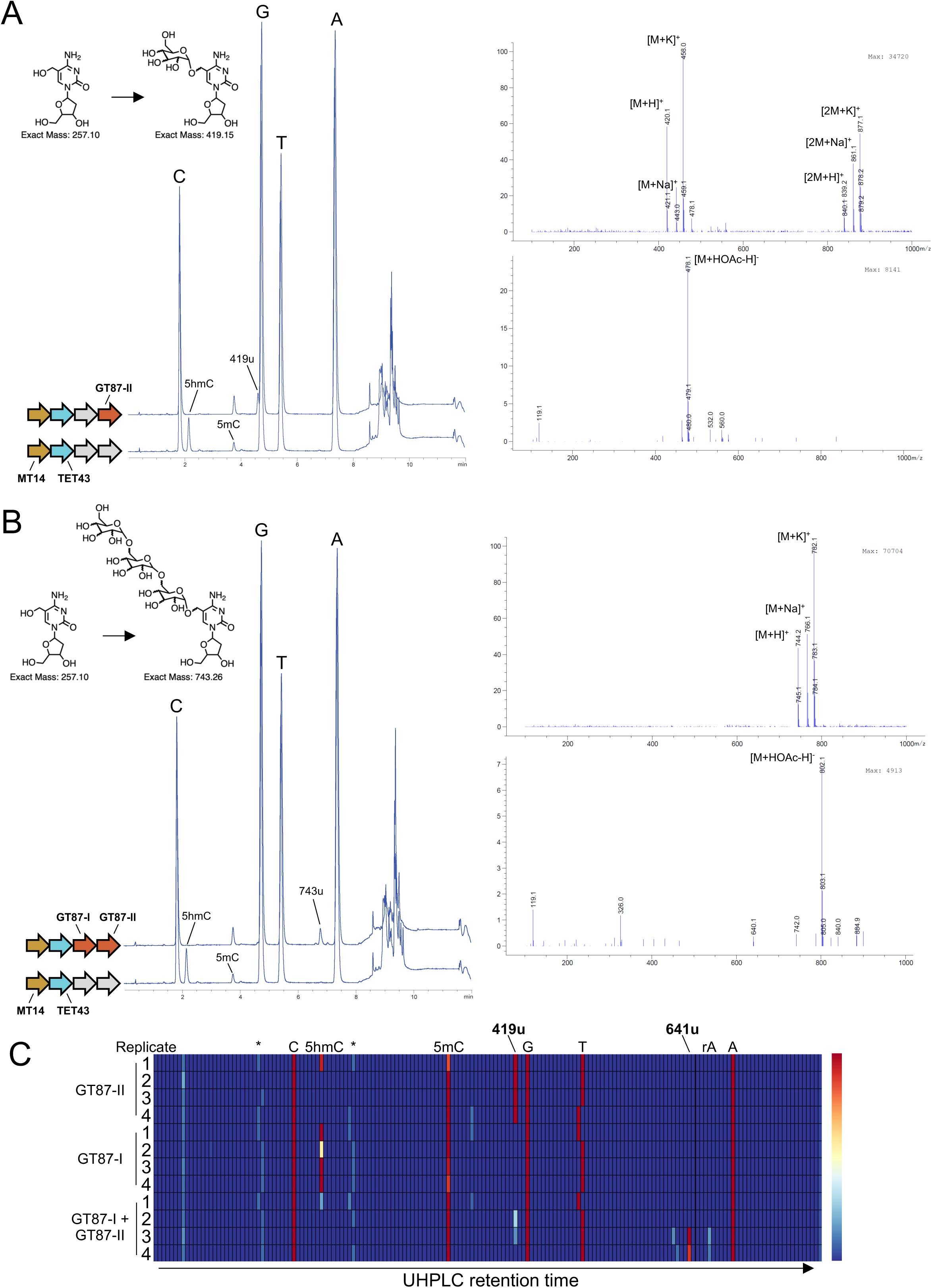
LC-MS analyses of new sugar modifications installed by subclass I GTs. (A) LC-MS analysis of nucleosides from gDNA isolated from *E. coli* expressing GT87-II which installs the first sugar on 5hmC precursors. LC trace (left) and MS spectra in positive (right, top) and negative (right, bottom) modes are shown. Expression vector assemblies are shown next to their corresponding LC traces. Empty grey arrows represent positions in the pSL126 vector assembled with non-coding pET28a control amplicons to control for enzyme position and expression plasmid size. MS spectra are shown for the new peak at retention time ∼4.58 min showing a mass of 419u. (B) LC-MS analysis of nucleosides from gDNA isolated from *E. coli* expressing GT87-I and GT87-II showing installation of subsequent sugars on 5hmC precursors. LC trace (left) and MS spectra in positive (right, top) and negative (right, bottom) modes are shown. Expression vector assemblies are shown next to their corresponding LC traces. Empty grey arrows represent positions in the pSL126 vector assembled with non-coding pET28a control amplicons to control for enzyme position and expression plasmid size. MS spectra are shown for the new peak at retention time ∼6.77 min showing a mass of 743u. (C) Heat map visualization of UHPLC results for contig 87 enzymes expressed in *E. coli*. Colored boxes represent peaks from the UHPLC traces. Heat map scale by color intensity is shown to the right and corresponds to total normalized area for peaks across retention time intervals. Results from four biological replicates are shown for each condition. The elution of non-specific and low-abundance product peaks of unknown masses are indicated with asterisks.

**Figure S4.**
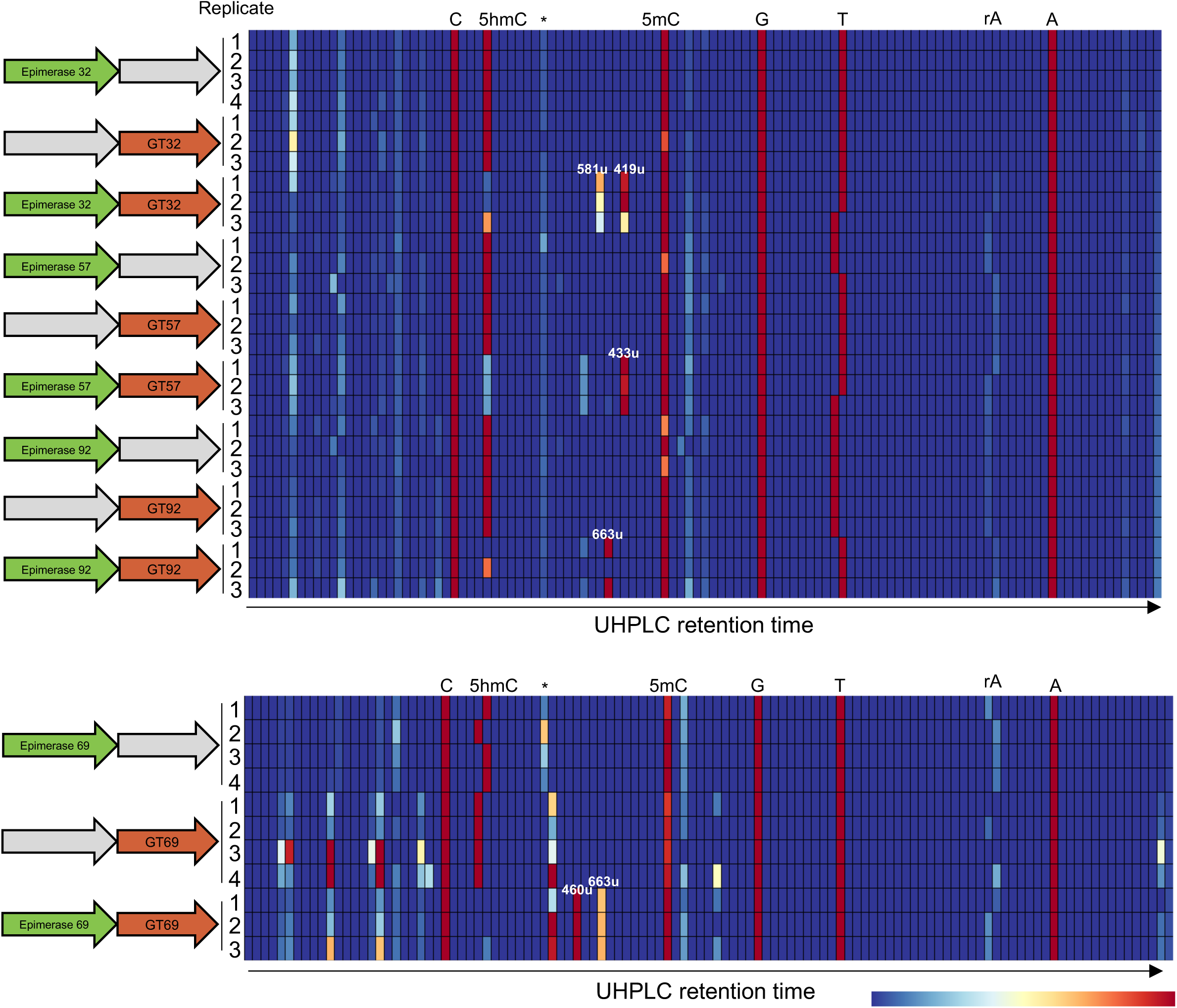
Analyses of new sugar modifications installed by subclass II GTs. Heat map visualization of total nucleosides from *E. coli* expressing subclass II GTs with their corresponding sugar epimerases. All expression constructs encode MT14 and TET43 for biosynthesis of 5hmC precursor residues. Empty grey arrows represent positions in the pSL126 vector assembled with non-coding pET28a control amplicons to control for enzyme position and expression plasmid size. Masses for new products are labeled where appropriate. Results from 3-4 biological replicates are shown for each condition. The elution of non-specific and low-abundance product peaks of unknown masses are indicated with asterisks.

**Figure S5.**
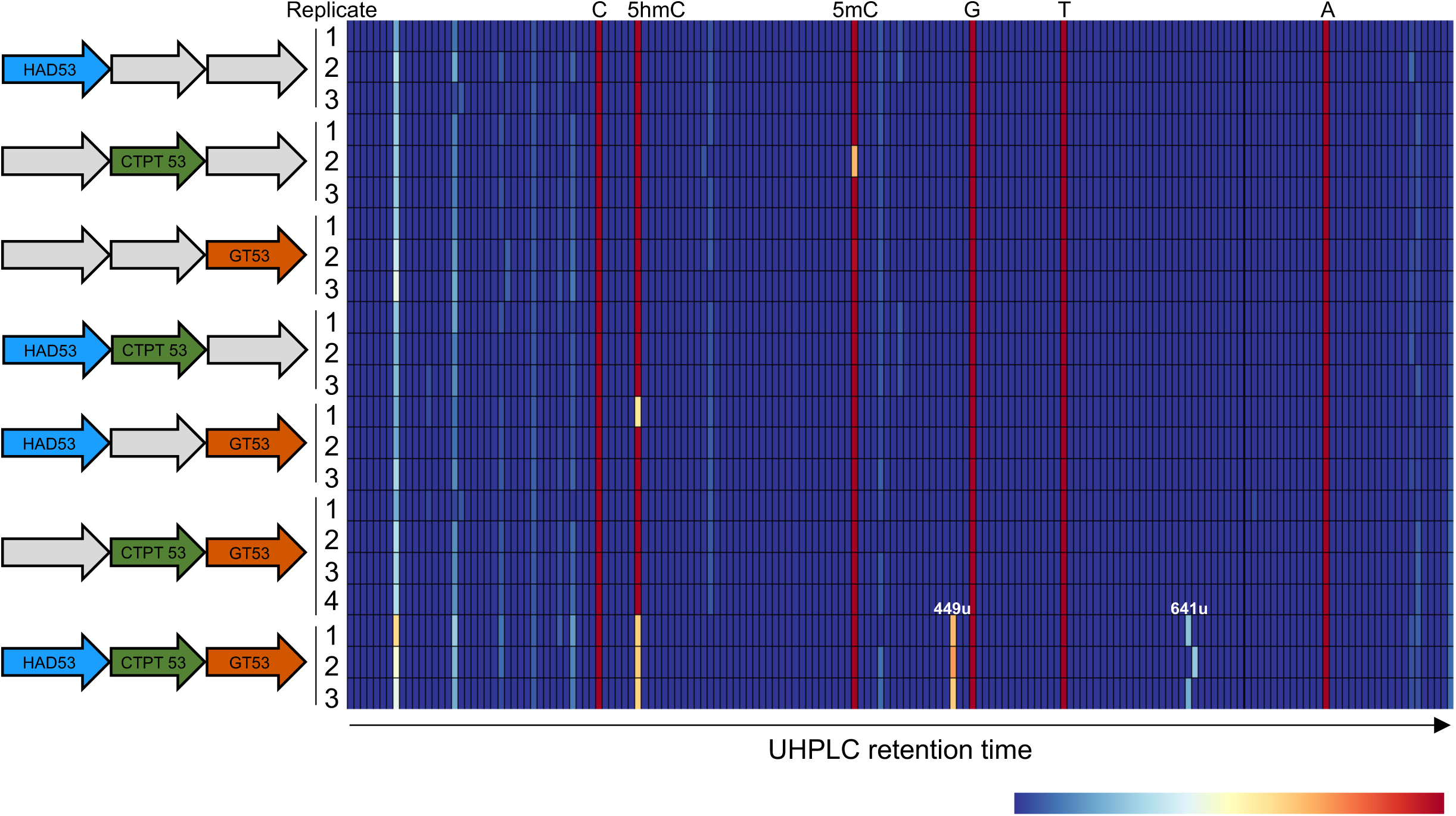
Analyses of new sugar modifications installed by contig 53 enzymes. Heat map visualization of total nucleosides from *E. coli* expressing enzymes from the contig 53 BGC. All expression constructs encode MT14 and TET43 for biosynthesis of 5hmC precursor residues. Empty grey arrows represent positions in the pSL126 vector assembled with non-coding pET28a control amplicons to control for enzyme position and expression plasmid size. Masses for new products are labeled where appropriate. Results from 3-4 biological replicates are shown for each condition.

**Figure S6.**
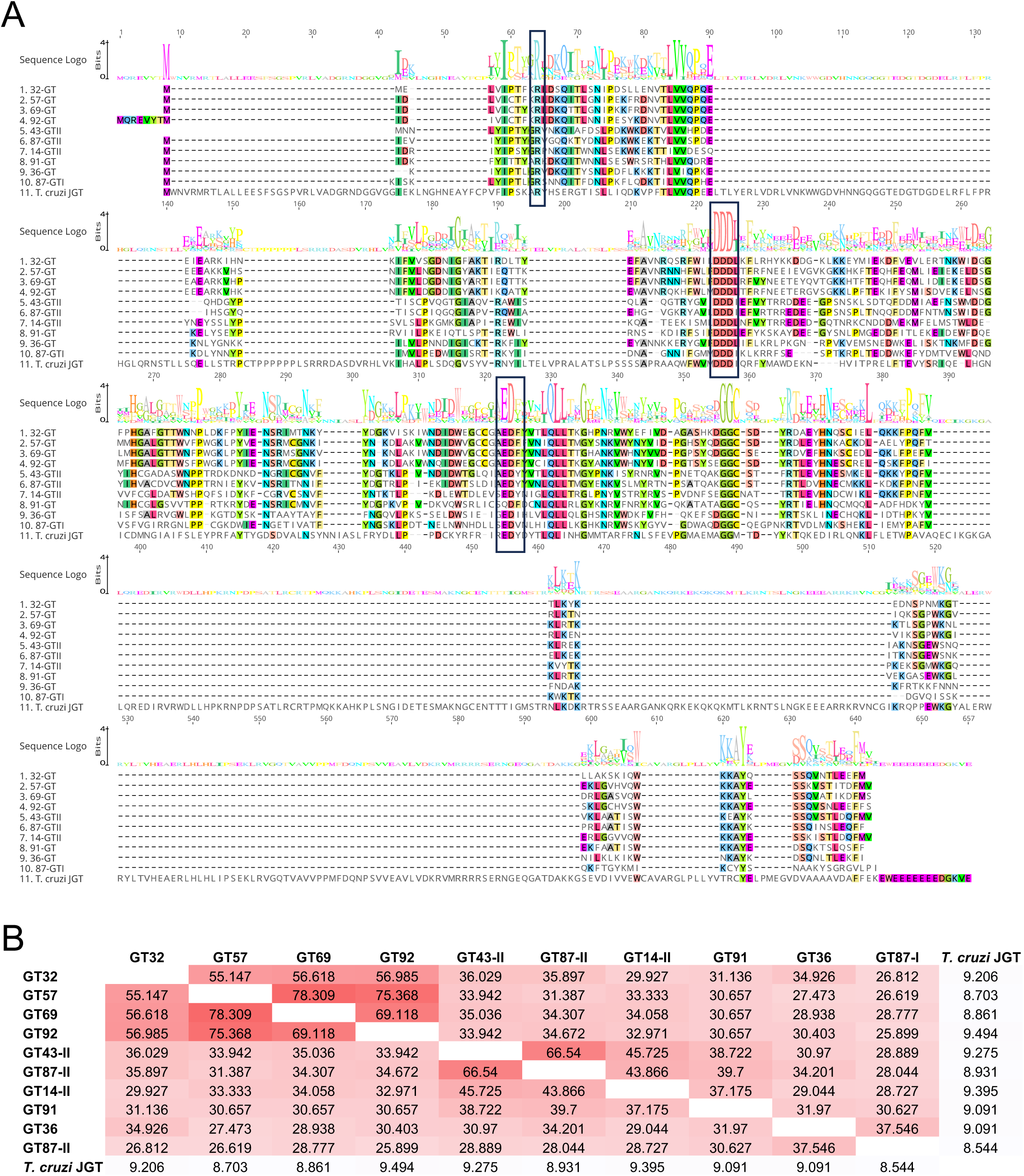
Alignment of catalytic site residues for phage GT-A enzymes and *T. cruzi* JGT. Multiple sequence alignments were generated as described in Materials and Methods. Conserved active site residues for the GT-A family are indicated with black boxes. (B) Pairwise analysis of amino acid sequence similarity for phage GT-A enzymes and *T. cruzi* JGT. Boxes are colored by increasing sequence similarity.

**Figure S7.**
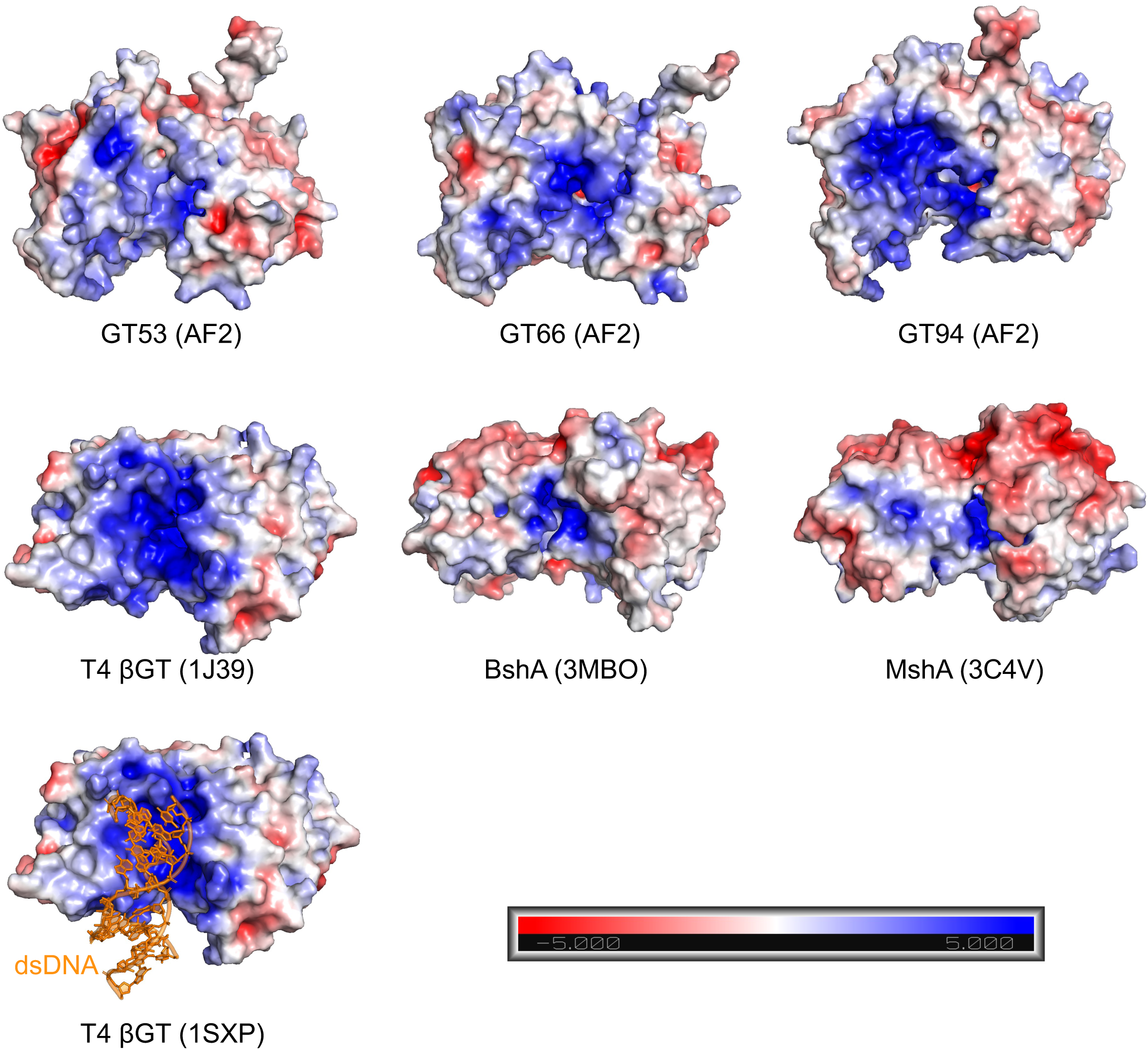
Electrostatic surface visualization of GT-B fold-containing phage GTs and top structural homologs. Structural models are positioned to highlight the positively-charged surface-exposed residues between the GT-B Rossmann folds. The T4 BGT structure is represented with and without bound double-stranded DNA (dsDNA). Scale shown represents relative charge potential of the surface-exposed residues.

**Figure S8.**
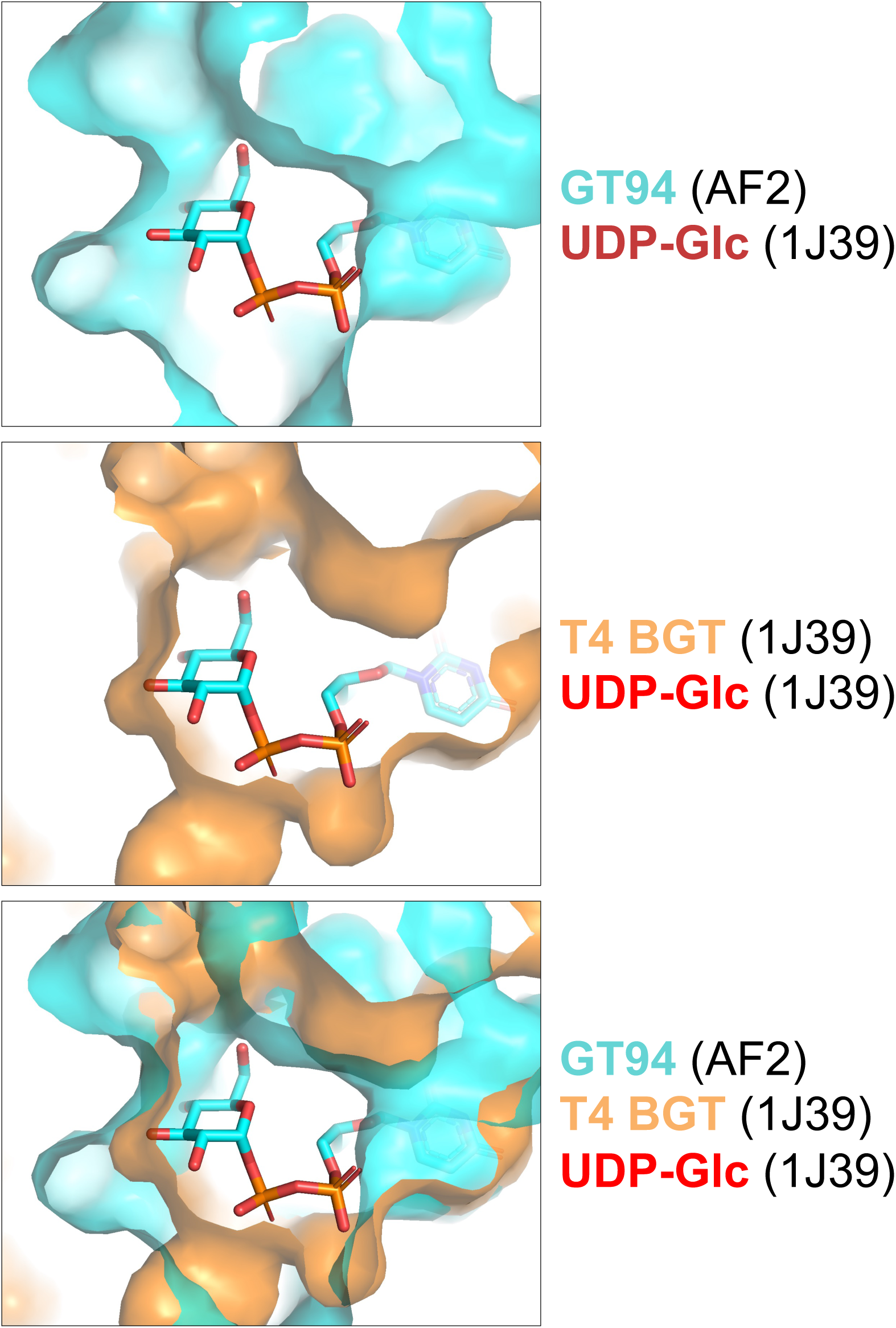
Surface representation of the putative sugar binding pocket of GT94 (top) compared to the structure of UDP-glucose (UDP-Glc)-bound T4 BGT (middle, 1J39). An overlay of the AlphaFold2 (AF2) GT94 structural model and the T4 BGT structure is shown to highlight differences in the solvent-exposed cavity surrounding the glucose moiety (bottom).

**Figure S9.**
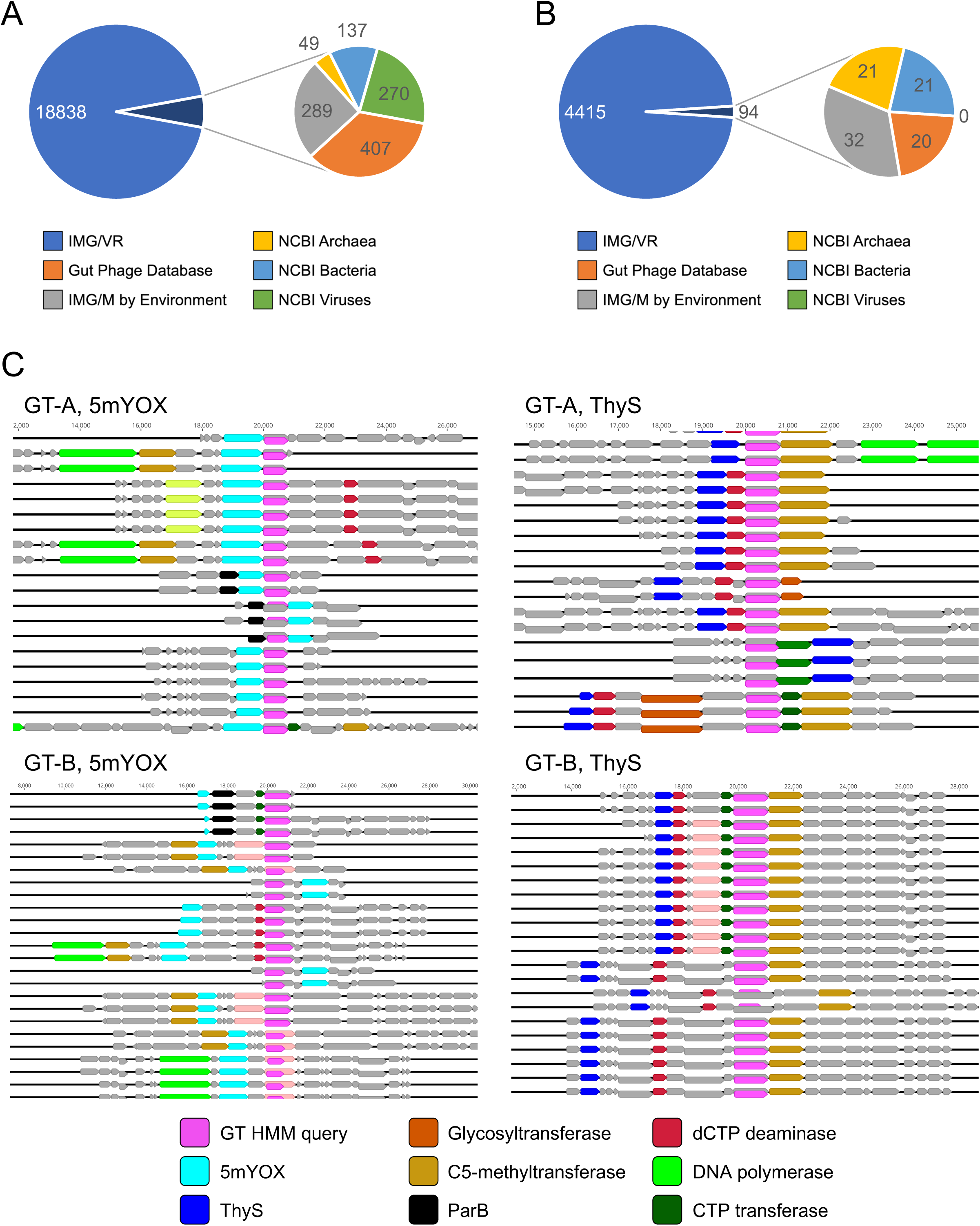
Summary of new GT-A (A) and GT-B (B) containing pathways identified through database mining using HMM profiles. Pie charts represent the total numbers of pathways identified by the different genomic and metagenomic databases probed. (C) Example contig architectures for GT-A- (top) and GT-B-containing (bottom) contigs paired with a 5mYOX (left) or ThyS (right) annotation. Genes containing additional domains of interest are colored according to the key shown at the bottom.

**Figure S10.**
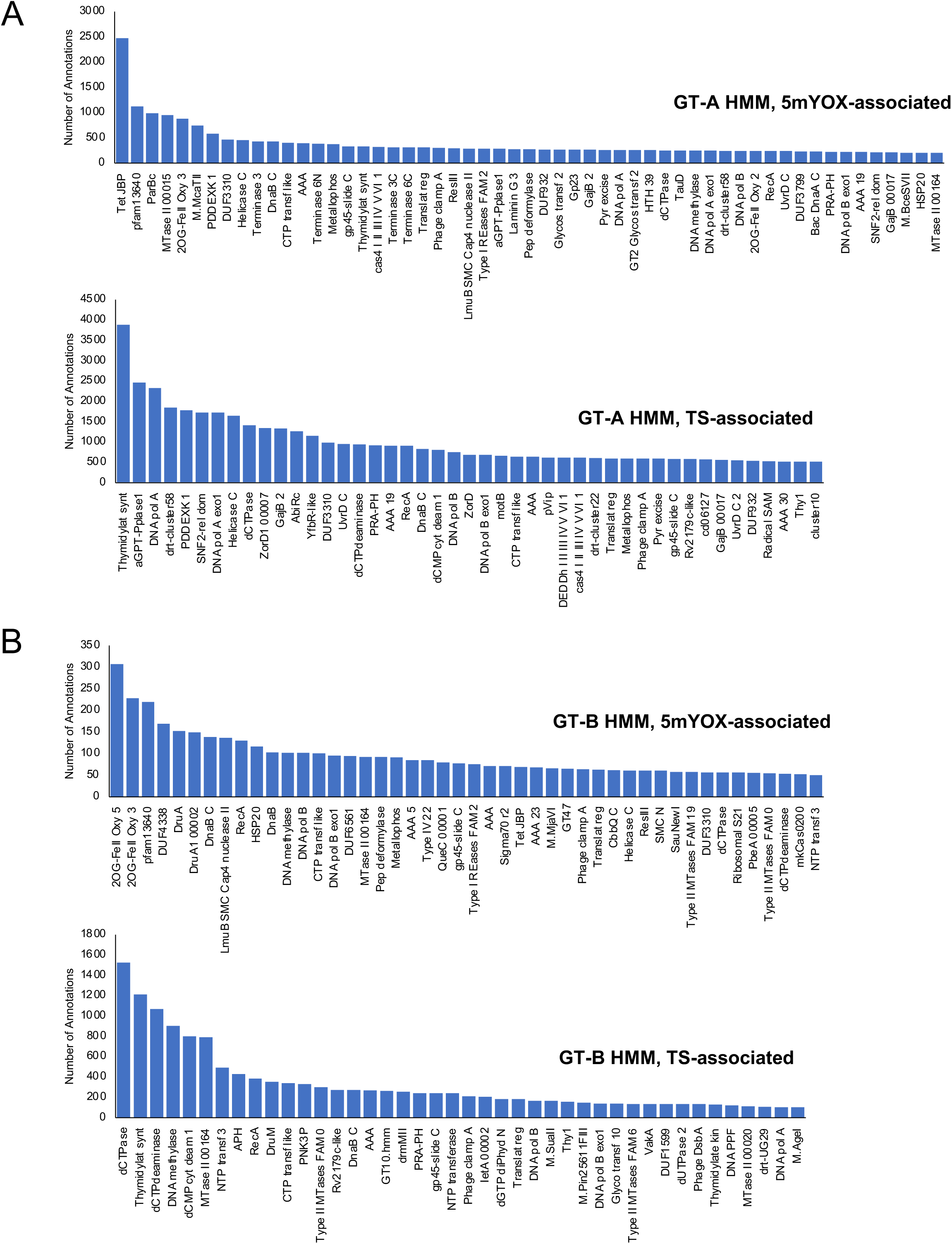
Frequency of gene annotations co-occurring in new GT-containing pathways. Contigs isolated from the GT-A (A) and GT-B (B) HMM profile searches were separated into 5mYOX-containing (A, B, top) or TS-containing (A, B, bottom) pathways. These groups containg a GT and 5mYOX or TS were further categorized by co-occurrence with additional domain annotations from Pfam, CAZy, DefenseFinder, and PADLOC. Bar graphs represent the total counts (y axis) of assigned annotations (x axis) per subgroup. All domains represented by more than n=250 (A) or n=50 (B) unique annotations in the mined dataset are included and presented from most to least frequent occurrence (left to right on graphs). Annotations in the datasets with fewer than n=250 (A) or n=50 (B) are not included for space limitations.

**Figure S11.**
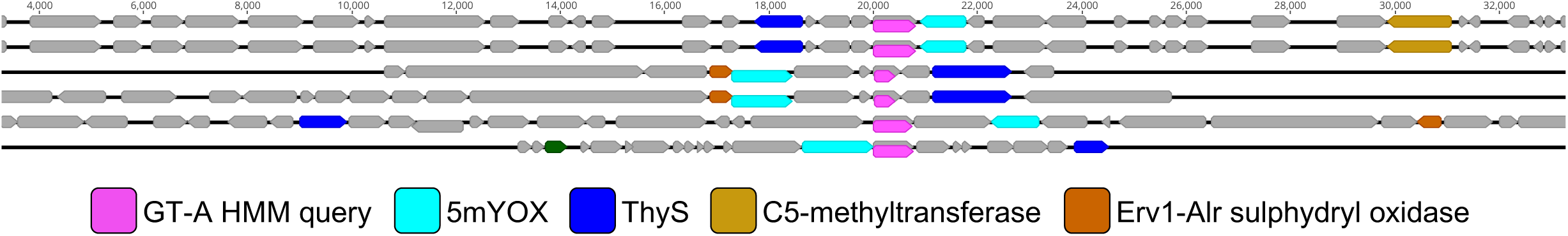
Six contigs containing genes with predicted 5mYOX and ThyS functions. Pathways are organized around the GT-A HMM query hit (magenta) with approximately 10 coding regions included on either side. Non-coding regions or contigs that are shorter than the 10 coding region cutoff length are represented by solid black lines. Coding regions of interest are colored according to frequently occurring domain annotations show in the key below.

**Figure S12.**
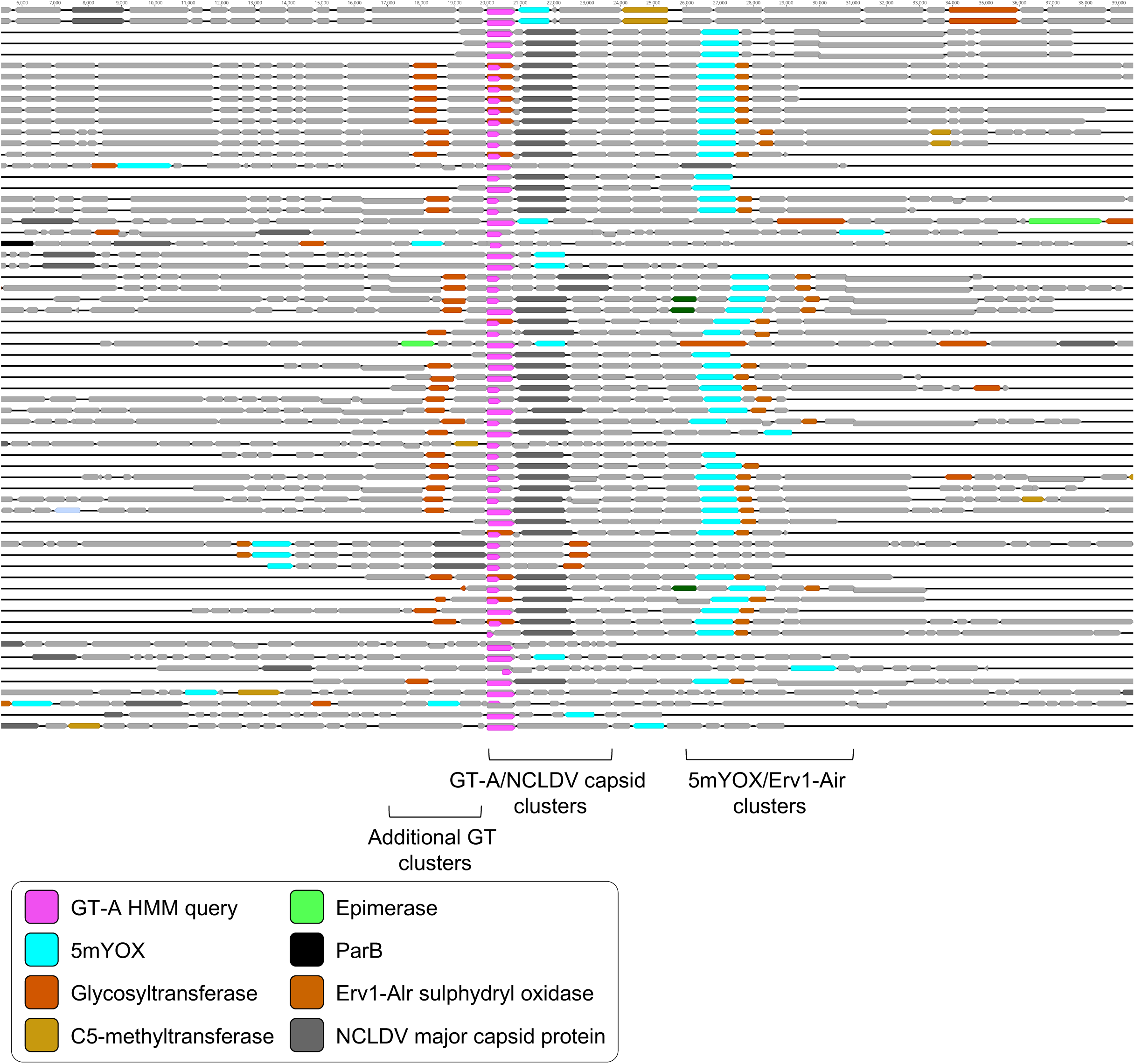
Illustration of GT-A, 5mYOX, and NCLDV capsid-containing metagenomic neighborhoods. Pathways are organized around the GT-A HMM query hit (magenta) with approximately 10 kb of additional sequence included on either side. Non-coding regions or contigs that are shorter than the ∼10 kb cutoff length are represented by solid black lines. Coding regions of interest are colored according to frequently occurring domain annotations show in the key below. Grey coding regions contain additional domains not highlighted for clarity or are uncharacterized.

**Figure S13.**
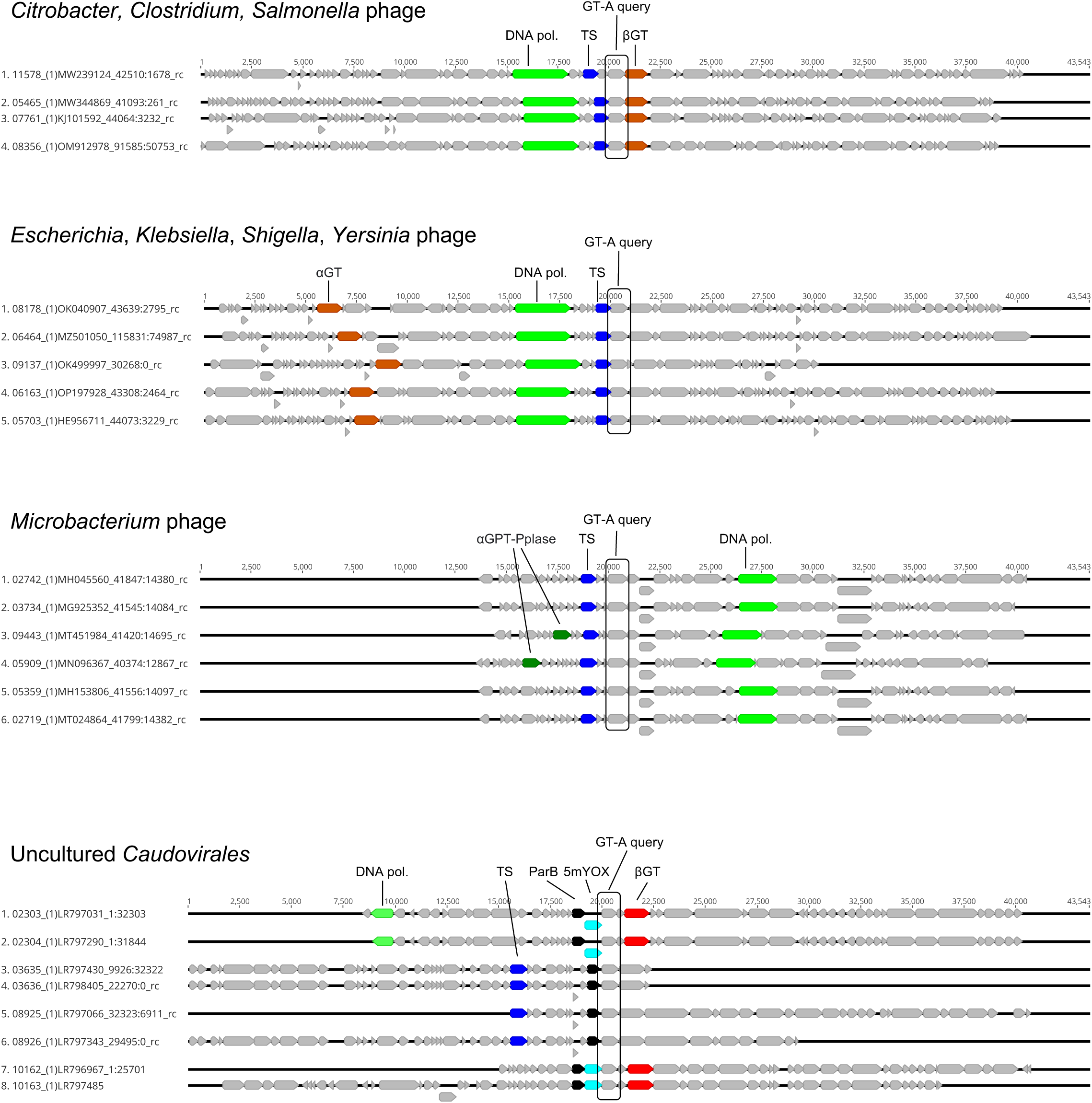
GT-A-like enzymes identified in sequenced viral genomes. Phage genomes in NCBI were identified as containing a GT based on sequence similarity from our GT-A HMM profile search. Representatives of the four major classes of phage taxa are shown. The complete list of genomes identified is included in Table S4. Relative positions of the GT-A query hit and other key genes of interest are shown for each group of phage. Pathways are organized around the GT-A HMM query hit with 20 kb of coding regions included on either side. Non-coding regions or genomes that are cut short of the 20 kb flanking region length are represented by solid black lines.

**Figure S14.**
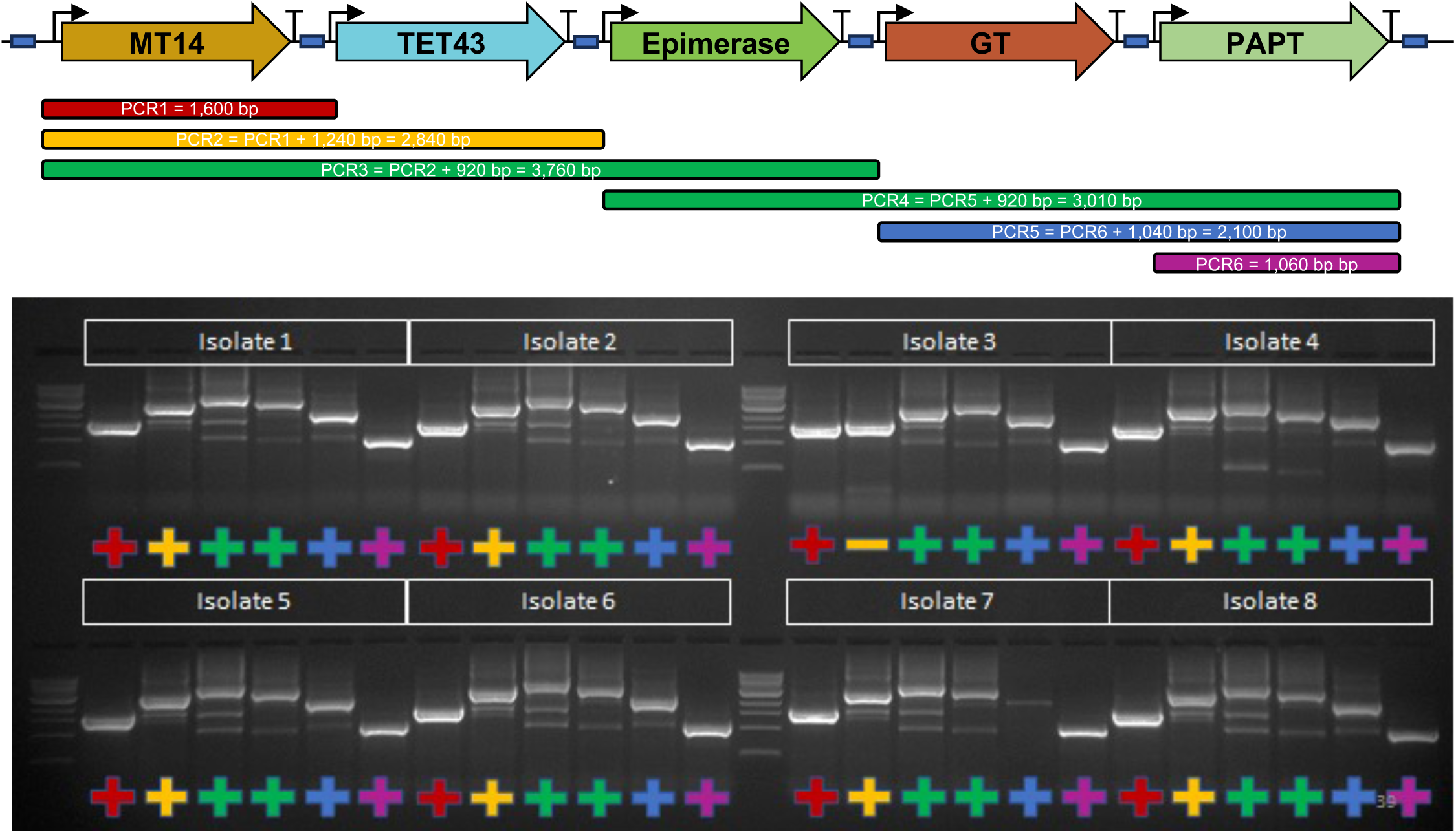
Example of contig 69 assembly verification by colony PCR (cPCR). Eight separate colonies were subjected to cPCR using primers specific to the intergenic regions flanking the GGA overlap sites in between genes (blue rectangle). Amplicon lengths are represented by colored boxes below the assembly illustration and correspond to amplicons shown on the ethidium bromide-stained gel below. PCRs that did not pass verification are indicated with a minus symbol “-“.

## REFERENCES

1. B. Macek, et al., Protein post-translational modifications in bacteria. Nat Rev Microbiol 17, 651–664 (2019).

2. A. Breiling, F. Lyko, Epigenetic regulatory functions of DNA modifications: 5-methylcytosine and beyond. Epigenetics Chromatin 8, 24 (2015).

3. W. V. Gilbert, T. A. Bell, C. Schaening, Messenger RNA modifications: Form, distribution, and function. Science (1979) 352, 1408–1412 (2016).

4. L. M. Iyer, S. Abhiman, L. Aravind, “Chapter 2 - Natural History of Eukaryotic DNA Methylation Systems” in Progress in Molecular Biology and Translational Science, X. Cheng, R. M. Blumenthal, Eds. (Academic Press, 2011), pp. 25–104.

5. E. V. Koonin, K. S. Makarova, Y. I. Wolf, Evolutionary Genomics of Defense Systems in Archaea and Bacteria. Annu Rev Microbiol 71, 233–261 (2017).

6. R. Pecori, S. Di Giorgio, J. Paulo Lorenzo, F. Nina Papavasiliou, Functions and consequences of AID/APOBEC-mediated DNA and RNA deamination. Nat Rev Genet 23, 505–518 (2022).

7. S. G. Conticello, The AID/APOBEC family of nucleic acid mutators. Genome Biol 9, 229 (2008).

8. R. W. Hendrix, M. C. M. Smith, R. N. Burns, M. E. Ford, G. F. Hatfull, Evolutionary relationships among diverse bacteriophages and prophages: All the world’s a phage. Proceedings of the National Academy of Sciences 96, 2192–2197 (1999).

9. A. R. Mushegian, Are There 1031 Virus Particles on Earth, or More, or Fewer? J Bacteriol 202, e00052–20 (2020).

10. E. V. Koonin, M. Krupovic, V. V. Dolja, The global virome: How much diversity and how many independent origins? Environ Microbiol 25, 40–44 (2023).

11. L. P. P. Braga, et al., Viruses direct carbon cycling in lake sediments under global change. Proceedings of the National Academy of Sciences 119, e2202261119 (2022).

12. R. J. Puxty, A. D. Millard, D. J. Evans, D. J. Scanlan, Viruses Inhibit CO2 Fixation in the Most Abundant Phototrophs on Earth. Current Biology 26, 1585–1589 (2016).

13. E. R. Westra, A. Buckling, P. C. Fineran, CRISPR–Cas systems: beyond adaptive immunity. Nat Rev Microbiol 12, 317–326 (2014).

14. E. V. Koonin, K. S. Makarova, CRISPR-Cas: Evolution of an RNA-based adaptive immunity system in prokaryotes. RNA Biol 10, 679–686 (2013).

15. J. H. Gommers-Ampt, P. Borst, Hypermodified bases in DNA. The FASEB Journal 9, 1034–1042 (1995).

16. Y.-J. Lee, et al., Pathways of thymidine hypermodification. Nucleic Acids Res 50, 3001– 3017 (2022).

17. P. R. Weigele, E. A. Raleigh, Biosynthesis and Function of Modified Bases in Bacteria and Their Viruses. Chem Rev 116, 12655–12687 (2016).

18. E. J. Burke, et al., Phage-encoded ten-eleven translocation dioxygenase (TET) is active in C5-cytosine hypermodification in DNA. Proceedings of the National Academy of Sciences 118, e2026742118 (2021).

19. Y.-J. Lee, et al., Identification and biosynthesis of thymidine hypermodifications in the genomic DNA of widespread bacterial viruses. Proceedings of the National Academy of Sciences 115, E3116–E3125 (2018).

20. R. M. Kohli, Y. Zhang, TET enzymes, TDG and the dynamics of DNA demethylation. Nature 502, 472–479 (2013).

21. X. Yin, L. Hu, Y. Xu, “Structure and Function of TET Enzymes” in DNA Methyltransferases - Role and Function, A. Jeltsch, R. Z. Jurkowska, Eds. (Springer International Publishing, 2022), pp. 239–267.

22. E. Tamanaha, S. Guan, K. Marks, L. Saleh, Distributive Processing by the Iron(II)/α-Ketoglutarate-Dependent Catalytic Domains of the TET Enzymes Is Consistent with Epigenetic Roles for Oxidized 5-Methylcytosine Bases. J Am Chem Soc 138, 9345–9348 (2016).

23. H. Hashimoto, et al., Structure of a Naegleria Tet-like dioxygenase in complex with 5-methylcytosine DNA. Nature 506, 391–395 (2014).

24. L. M. Iyer, D. Zhang, A. Maxwell Burroughs, L. Aravind, Computational identification of novel biochemical systems involved in oxidation, glycosylation and other complex modifications of bases in DNA. Nucleic Acids Res 41, 7635–7655 (2013).

25. H. Geoffrey, L. Yan-Jiun, de C.-L. Valérie, P. R. Weigele, Hypermodified DNA in Viruses of E. coli and Salmonella. EcoSal Plus 9, eESP-0028-2019 (2021).

26. I. R. Lehman, E. A. Pratt, On the Structure of the Glucosylated Hydroxymethylcytosine Nucleotides of Coliphages T2, T4, and T6. Journal of Biological Chemistry 235, 3254–3259 (1960).

27. J. L. Munro, J. S. Wiberg, Evidence that gene 56 of bacteriophage T4 is a structural gene for deoxycytidine triphosphatase. Virology 36, 442–446 (1968).

28. S. R. Kornberg, S. B. Zimmerman, A. Kornberg, Glucosylation of Deoxyribonucleic Acid by Enzymes from Bacteriophage-infected Escherichia coli. Journal of Biological Chemistry 236, 1487–1493 (1961).

29. L. M. Gold, M. Schweiger, Synthesis of phage-specific α- and β-glucosyl transferases directed by T-even DNA in vitro. Proceedings of the National Academy of Sciences 62, 892–898 (1969).

30. K. Flodman, et al., Type II Restriction of Bacteriophage DNA With 5hmdU-Derived Base Modifications. Front Microbiol 10 (2019).

31. L.-H. Huang, C. M. Farnet, K. C. Ehrlich, M. Ehrlich, Digestion of highly modified bacteriophage DNA by restriction endonucleases. Nucleic Acids Res 10, 1579–1591 (1982).

32. P. B. Miller, W. W. Wakarchuk, R. A. J. Warren, α -Putrescinylthymine and the sensitivity of bacteriophage φ W-14 DNA to restriction endonucleases. Nucleic Acids Res 13, 2559–2568 (1985).

33. R. Tsai, I. R. Corrêa, M. Y. Xu, S. Xu, Restriction and modification of deoxyarchaeosine (dG+)-containing phage 9 g DNA. Sci Rep 7, 8348 (2017).

34. D. H. Krüger, T. A. Bickle, Bacteriophage survival: multiple mechanisms for avoiding the deoxyribonucleic acid restriction systems of their hosts. Microbiol Rev 47, 345–360 (1983).

35. D. Paez-Espino, et al., IMG/VR: a database of cultured and uncultured DNA Viruses and retroviruses. Nucleic Acids Res 45, gkw1030 (2017).

36. A. C. Gregory, et al., Marine DNA Viral Macro- and Microdiversity from Pole to Pole. Cell 177, 1109–1123.e14 (2019).

37. C. Engler, R. Gruetzner, R. Kandzia, S. Marillonnet, Golden Gate Shuffling: A One-Pot DNA Shuffling Method Based on Type IIs Restriction Enzymes. PLoS One 4, e5553-(2009).

38. V. Potapov, et al., Comprehensive Profiling of Four Base Overhang Ligation Fidelity by T4 DNA Ligase and Application to DNA Assembly. ACS Synth Biol 7, 2665–2674 (2018).

39. J. M. Pryor, et al., Enabling one-pot Golden Gate assemblies of unprecedented complexity using data-optimized assembly design. PLoS One 15, e0238592- (2020).

40. K. A. Padgett, J. A. Sorge, Creating seamless junctions independent of restriction sites in PCR cloning. Gene 168, 31–35 (1996).

41. J. M. Pryor, V. Potapov, K. Bilotti, N. Pokhrel, G. J. S. Lohman, Rapid 40 kb Genome Construction from 52 Parts through Data-optimized Assembly Design. ACS Synth Biol 11, 2036–2042 (2022).

42. J. Jumper, et al., Highly accurate protein structure prediction with AlphaFold. Nature 596, 583–589 (2021).

43. M. van Kempen, et al., Fast and accurate protein structure search with Foldseek. Nat Biotechnol (2023) 10.1038/s41587-023-01773-0.

44. The UniProt Consortium, UniProt: the Universal Protein Knowledgebase in 2023. Nucleic Acids Res 51, D523–D531 (2023).

45. W. Bullard, J. Lopes da Rosa-Spiegler, S. Liu, Y. Wang, R. Sabatini, Identification of the Glucosyltransferase That Converts Hydroxymethyluracil to Base J in the Trypanosomatid Genome *. Journal of Biological Chemistry 289, 20273–20282 (2014).

46. D. Gerlach, et al., Methicillin-resistant Staphylococcus aureus alters cell wall glycosylation to evade immunity. Nature 563, 705–709 (2018).

47. D. Gerlach, et al., Horizontal transfer and phylogenetic distribution of the immune evasion factor tarP. Front Microbiol 13 (2022).

48. X. Guoqing, et al., Wall Teichoic Acid-Dependent Adsorption of Staphylococcal Siphovirus and Myovirus. J Bacteriol 193, 4006–4009 (2011).

49. C. A. Schnaitman, J. D. Klena, Genetics of lipopolysaccharide biosynthesis in enteric bacteria. Microbiol Rev 57, 655–682 (1993).

50. E. Pradel, C. T. Parker, C. A. Schnaitman, Structures of the rfaB, rfaI, rfaJ, and rfaS genes of Escherichia coli K-12 and their roles in assembly of the lipopolysaccharide core. J Bacteriol 174, 4736–4745 (1992).

51. C. W. Price, et al., Genome-wide analysis of the general stress response in Bacillus subtilis. Mol Microbiol 41, 757–774 (2001).

52. I.-M. A. Chen, et al., The IMG/M data management and analysis system v.7: content updates and new features. Nucleic Acids Res 51, D723–D732 (2023).

53. L. F. Camarillo-Guerrero, A. Almeida, G. Rangel-Pineros, R. D. Finn, T. D. Lawley, Massive expansion of human gut bacteriophage diversity. Cell 184, 1098–1109.e9 (2021).

54. U. Neri, et al., Expansion of the global RNA virome reveals diverse clades of bacteriophages. Cell 185, 4023–4037.e18 (2022).

55. J. Mistry, et al., Pfam: The protein families database in 2021. Nucleic Acids Res 49, D412–D419 (2021).

56. D. H. Haft, et al., TIGRFAMs: a protein family resource for the functional identification of proteins. Nucleic Acids Res 29, 41–43 (2001).

57. R. J. Roberts, T. Vincze, J. Posfai, D. Macelis, REBASE: a database for DNA restriction and modification: enzymes, genes and genomes. Nucleic Acids Res 51, D629–D630 (2023).

58. E. Drula, et al., The carbohydrate-active enzyme database: functions and literature. Nucleic Acids Res 50, D571–D577 (2022).

59. L. J. Payne, et al., PADLOC: a web server for the identification of antiviral defence systems in microbial genomes. Nucleic Acids Res 50, W541–W550 (2022).

60. F. Tesson, et al., Systematic and quantitative view of the antiviral arsenal of prokaryotes. Nat Commun 13, 2561 (2022).

61. F. O. Aylward, M. Moniruzzaman, A. D. Ha, E. V. Koonin, A phylogenomic framework for charting the diversity and evolution of giant viruses. PLoS Biol 19, e3001430- (2021).

62. A. Chakrapani, et al., Glucosylated 5-Hydroxymethylpyrimidines as Epigenetic DNA Bases Regulating Transcription and Restriction Cleavage. Chemistry (Easton) 28, e202200911 (2022).

63. R. S. Ayikpoe, et al., A scalable platform to discover antimicrobials of ribosomal origin. Nat Commun 13, 6135 (2022).

64. V. R. Chaudhari, M. R. Hanson, GoldBricks: an improved cloning strategy that combines features of Golden Gate and BioBricks for better efficiency and usability. Synth Biol 6, ysab032 (2021).

65. F. Yan, et al., Synthetic biology approaches and combinatorial biosynthesis towards heterologous lipopeptide production. Chem Sci 9, 7510–7519 (2018).

66. H. Ren, P. Hu, H. Zhao, A plug-and-play pathway refactoring workflow for natural product research in Escherichia coli and Saccharomyces cerevisiae. Biotechnol Bioeng 114, 1847–1854 (2017).

67. D. Chiasson, et al., A unified multi-kingdom Golden Gate cloning platform. Sci Rep 9, 10131 (2019).

68. Y. Tong, J. Zhou, L. Zhang, P. Xu, A Golden-Gate Based Cloning Toolkit to Build Violacein Pathway Libraries in Yarrowia lipolytica. ACS Synth Biol 10, 115–124 (2021).

69. N. K. Duggal, M. Emerman, Evolutionary conflicts between viruses and restriction factors shape immunity. Nat Rev Immunol 12, 687–695 (2012).

70. M. Krupovic, V. V. Dolja, E. V. Koonin, The virome of the last eukaryotic common ancestor and eukaryogenesis. Nat Microbiol 8, 1008–1017 (2023).

71. J. Lombard, The multiple evolutionary origins of the eukaryotic N-glycosylation pathway. Biol Direct 11, 36 (2016).

72. R. Taujale, et al., Deep evolutionary analysis reveals the design principles of fold A glycosyltransferases. Elife 9, e54532 (2020).

73. M. A. Katz, et al., Diverse viral cas genes antagonize CRISPR immunity. bioRxiv, 2023.06.24.545427 (2023).

74. F. Rousset, et al., Phages and their satellites encode hotspots of antiviral systems. Cell Host Microbe 30, 740–753.e5 (2022).

75. M. G. Fischer, T. Hackl, Host genome integration and giant virus-induced reactivation of the virophage mavirus. Nature 540, 288–291 (2016).

76. T. Hackl, S. Duponchel, K. Barenhoff, A. Weinmann, M. G. Fischer, Virophages and retrotransposons colonize the genomes of a heterotrophic flagellate. Elife 10, e72674 (2021).

77. C. Peipei, et al., LamB, OmpC, and the Core Lipopolysaccharide of Escherichia coli K-12 Function as Receptors of Bacteriophage Bp7. J Virol 94, e00325–20 (2020).

78. J. Bertozzi Silva, Z. Storms, D. Sauvageau, Host receptors for bacteriophage adsorption. FEMS Microbiol Lett 363, fnw002 (2016).

79. R. Sandulache, P. Prehm, D. Kamp, Cell wall receptor for bacteriophage Mu G(+). J Bacteriol 160, 299–303 (1984).

80. S. Yokota, T. Hayashi, H. Matsumoto, Identification of the lipopolysaccharide core region as the receptor site for a cytotoxin-converting phage, phi CTX, of Pseudomonas aeruginosa. J Bacteriol 176, 5262–5269 (1994).

81. A. A. Filippov, et al., Bacteriophage-Resistant Mutants in Yersinia pestis: Identification of Phage Receptors and Attenuation for Mice. PLoS One 6, e25486- (2011).

82. K. E. Kortright, B. K. Chan, P. E. Turner, High-throughput discovery of phage receptors using transposon insertion sequencing of bacteria. Proceedings of the National Academy of Sciences 117, 18670–18679 (2020).

83. J. L. Kadrmas, C. R. H. Raetz, Enzymatic Synthesis of Lipopolysaccharide in *Escherichia coli*: PURIFICATION AND PROPERTIES OF HEPTOSYLTRANSFERASE I *. Journal of Biological Chemistry 273, 2799–2807 (1998).

84. J. Bondy-Denomy, et al., Prophages mediate defense against phage infection through diverse mechanisms. ISME J 10, 2854–2866 (2016).

85. K. Boeneman, E. Crooke, Chromosomal replication and the cell membrane. Curr Opin Microbiol 8, 143–148 (2005).

86. D. M. Knipe, A. Prichard, S. Sharma, J. Pogliano, Replication Compartments of Eukaryotic and Bacterial DNA Viruses: Common Themes Between Different Domains of Host Cells. Annu Rev Virol 9, 307–327 (2022).

87. E. G. Armbruster, et al., Sequential membrane- and protein-bound organelles compartmentalize genomes during phage infection. bioRxiv, 2023.09.20.558163 (2023).

88. J. Klumpp, D. E. Fouts, S. Sozhamannan, Next generation sequencing technologies and the changing landscape of phage genomics. Bacteriophage 2, 190–199 (2012).

89. J. Klumpp, et al., The odd one out: Bacillus ACT bacteriophage CP-51 exhibits unusual properties compared to related Spounavirinae W.Ph. and Bastille. Virology 462–463, 299–308 (2014).

90. G. Wolff, C. E. Melia, E. J. Snijder, M. Bárcena, Double-Membrane Vesicles as Platforms for Viral Replication. Trends Microbiol 28, 1022–1033 (2020).

91. T. G. Laughlin, et al., Architecture and self-assembly of the jumbo bacteriophage nuclear shell. Nature 608, 429–435 (2022).

92. E.-A. Raiber, et al., 5-Formylcytosine alters the structure of the DNA double helix. Nat Struct Mol Biol 22, 44–49 (2015).

93. M. Storch, et al., BASIC: A New Biopart Assembly Standard for Idempotent Cloning Provides Accurate, Single-Tier DNA Assembly for Synthetic Biology. ACS Synth Biol 4, 781–787 (2015).

94. Y.-J. Lee, P. R. Weigele, “Detection of Modified Bases in Bacteriophage Genomic DNA” in DNA Modifications: Methods and Protocols, A. Ruzov, M. Gering, Eds. (Springer US, 2021), pp. 53–66.

95. M. N. Price, P. S. Dehal, A. P. Arkin, FastTree 2 – Approximately Maximum-Likelihood Trees for Large Alignments. PLoS One 5, e9490- (2010).

96. C. L. M. Gilchrist, Y.-H. Chooi, clinker & clustermap.js: automatic generation of gene cluster comparison figures. Bioinformatics 37, 2473–2475 (2021).

97. L. Zimmermann, et al., A Completely Reimplemented MPI Bioinformatics Toolkit with a New HHpred Server at its Core. J Mol Biol 430, 2237–2243 (2018).

98. M. Mirdita, et al., ColabFold: making protein folding accessible to all. Nat Methods 19, 679–682 (2022).

99. S. R. Eddy, Accelerated Profile HMM Searches. PLoS Comput Biol 7, e1002195- (2011).

100. J. Liu, A. Mushegian, Three monophyletic superfamilies account for the majority of the known glycosyltransferases. Protein Science 12, 1418–1431 (2003).

101. L. Larivière, V. Gueguen-Chaignon, S. Moréra, Crystal Structures of the T4 Phage β-Glucosyltransferase and the D100A Mutant in Complex with UDP-glucose: Glucose Binding and Identification of the Catalytic Base for a Direct Displacement Mechanism. J Mol Biol 330, 1077–1086 (2003).

